# Adaptive duplication and functional diversification of Protein kinase R contribute to the uniqueness of bat-virus interactions

**DOI:** 10.1101/2022.06.28.497829

**Authors:** Stéphanie Jacquet, Michelle Culbertson, Chi Zang, Adil El Filali, Clément De La Myre Mory, Jean-Baptiste Pons, Ondine Filippi-Codaccioni, M. Elise Lauterbur, Barthélémy Ngoubangoye, Jeanne Duhayer, Clément Verez, Chorong Park, Clara Dahoui, Clayton M. Carey, Greg Brennan, David Enard, Andrea Cimarelli, Stefan Rothenburg, Nels C. Elde, Dominique Pontier, Lucie Etienne

## Abstract

Several bat species act as asymptomatic reservoirs for many viruses that are instead highly pathogenic in other mammals. Here, we have characterized the functional diversification of the Protein kinase R (PKR), a major antiviral innate defense system. Our data indicate that PKR has evolved under positive selection and has undergone repeated genomic duplications in bats, in contrast to all studied mammals that possess a single copy of the gene. Functional testing of the relationship between PKR and poxvirus antagonists revealed how an evolutionary conflict with ancient pathogenic poxviruses has shaped a specific bat host-virus interface. More importantly, we determined that duplicated PKRs of the *Myotis* species have undergone functional diversification allowing them to collectively escape from and enhance control of DNA and RNA viruses. These findings suggest that viral-driven adaptations in PKR contribute to modern virus-bat interactions and may account for bat specific immunity.

## Introduction

The present architecture of host innate immunity is the result of long-standing conflicts of ancient pathogenic viruses that continually adapted, and counter-adapted, to defeat or evade the antiviral defense of their host^1^. Hallmarks of these virus-host conflicts are the disproportionate accumulation of non-synonymous mutations and genetic novelties over evolutionary times at the interface of host antiviral effectors and viruses. While being the result of past viral pressure, these adaptations may explain why hosts are susceptible – or resistant – to modern-day viruses, and may also enlighten the functional diversity of host antiviral defenses^1,2^. Therefore, comparative functional genomics of hosts and viruses are of utmost importance to better understand what drives the specificity of virus-host interactions, particularly in wild host reservoirs of zoonotic viral pathogens^3^.

As the second most diverse and geographically widespread mammalian order, bats are outstanding among mammals because of their unique capability of powered flight and their propensity to a significant viral richness^4,5^. Several bat species are natural reservoirs for viruses that are highly virulent in other mammals, such as Marburg virus, Nipah virus, and SARS coronaviruses, without themselves showing symptoms^6^. These differences between bats and other mammals, in particular humans and non-human primates, have recently gathered considerable efforts to characterize the antiviral mechanisms of these flying mammals^3^. Bats may have evolved unique adaptations in their inflammasome components and signaling factors (*e.g*. PYHIN^7^, STING^8^, IRF3^9^, RIPK1^10^) that mitigate flight’s detrimental effects and dampen excessive inflammation, thereby presumably increasing viral tolerance. Furthermore, with more than 1,200 species and approximately 60 million years of divergence^11^, bats have co-evolved with a large diversity of viral pathogens. As a result, specific adaptive changes may also enable bats to efficiently control viral infections^12^. For example, a handful of bat antiviral factors bear signatures of strong positive selection and gene duplications^12,13^, including key restriction factors, such as APOBECs^12,14^, MX family GTPases^15^, IFITM3^16^ and TRIM5/22^17^. Nevertheless, efforts to broadly and comprehensively characterize the functional diversification of bat restriction factors, compared to other mammals, remain very limited. In particular, conclusions from most functional studies on bat immunity are primarily drawn from a specific bat species and a virus system. In-depth comparative and functional studies of bat antiviral effectors based on representative species are thus needed to decipher the diversity and specificities of chiropteran antiviral immune mechanisms.

Among the innate antiviral mechanisms, activation of the protein kinase R (PKR) constitutes one of the first line of mammalian antiviral defense. PKR is a keystone immune sensor and a broad restriction factor of a multitude of DNA and RNA viral families, such as Poxviridae, Herpesviridae and Orthomyxoviridae. Upon sensing of viral double-stranded RNA (dsRNA), PKR phosphorylates the alpha subunit of eukaryotic initiation factor 2α (eIF2α), leading to a potent cap-dependent translational shut down and viral inhibition. The importance of PKR in immunity is further highlighted by the fact that viruses have, in turn, developed various and strong antagonism mechanisms to circumvent PKR function^18–22^. One remarkable example is the mimicry of eIF2α by the poxvirus antagonist protein K3, which directly interacts with PKR to block eIF2α phosphorylation^23^. Over evolutionary times, PKR has continually been under pathogen’s pressure, as exemplified by its rapid adaptive evolution in primates and rabbits^19,24,25^.

In bats, how PKR has genetically and functionally evolved and how its past diversification contributes to modern bat immunity-virus interplays remain unknown. Here, we report deep functional adaptive changes and exceptional gene duplications in bat PKR that broaden escape mechanisms to viral antagonism, including from poxviruses, orthomyxoviruses, and herpesviruses, and enhance viral control in *Myotis* bats. Using an evolutionary-guided functional approach, we show that long-standing genetic conflicts with viral pathogens have driven the rapid evolution and duplication of bat PKRs, and the resulting adaptive changes account for modern host-virus antagonism specificity.

## Results

### PKR has been the target of strong diversifying selection and unusual gene duplication events in bats

The scarcity of bat genome sequences limits the study of their immunity, their virus-host interface and evolutionary history. To increase the robustness of our evolutionary analyses of bat PKRs, we sequenced and cloned additional coding sequences from 15 new bat species (see Materials and Methods). Overall, 33 bat orthologous sequences of PKR have been included in our positive selection analyses, spanning 62 million years of evolution ^11,26^. We compared models that disallow positive selection (models M1 and M7) to those allowing for positive selection (M2 and M8) using the PAML Codeml package^27^. We found that PKR has evolved under strong and recurrent positive selection during bat evolution, leading to significant adaptive signatures at both the gene and the codon levels (PAML codeml M1 *vs* M2 and M7 *vs* M8 p-values = 4.4 × 10^-83^ and 7.7 × 10^-86^, respectively Table S1).

To determine whether this adaptive evolution is common to all mammals, we extended our analysis to the four other major mammalian orders: Primata, Rodentia, Artiodactyla and Carnivora. We showed that rapid and recurrent evolution of PKR is common, with significant evidence of positive selection in all tested mammals (PAML codeml M1 *vs* M2 and M7 *vs* M8 p-values < 4.2 × 10^-4^ and 2.5 × 10^-5^, respectively Table S1). However, comparative analyses suggest more frequent signatures of adaptive changes in bat PKR, as well as marked differences in the location of the evolutionary footprints comparing to other orders. While most of the rapidly evolving sites are concentrated in the kinase domain of primate, artiodactyl and rodent PKRs (Figure 1A), the fast-evolving codons are scattered across bat PKR, with three remarkable hotspots in the second dsRNA binding-domain, the linker region, and the kinase domain (Figure 1A). Because bat lineages may have evolved under different selective pressures, we used a branch specific model (aBSREL) to test for episodic positive selection in PKR during bat evolution. We found that several bat lineages have been the targets of intensive episodic positive selection, in particular in the Yangochiroptera infra-order (Figure 1B), indicating differential pressure during bat evolution. Extending the branch analysis to other mammals showed that bat PKRs were among the most important targets of episodic positive selection across mammals (Figure S1).

**Figure 1.**
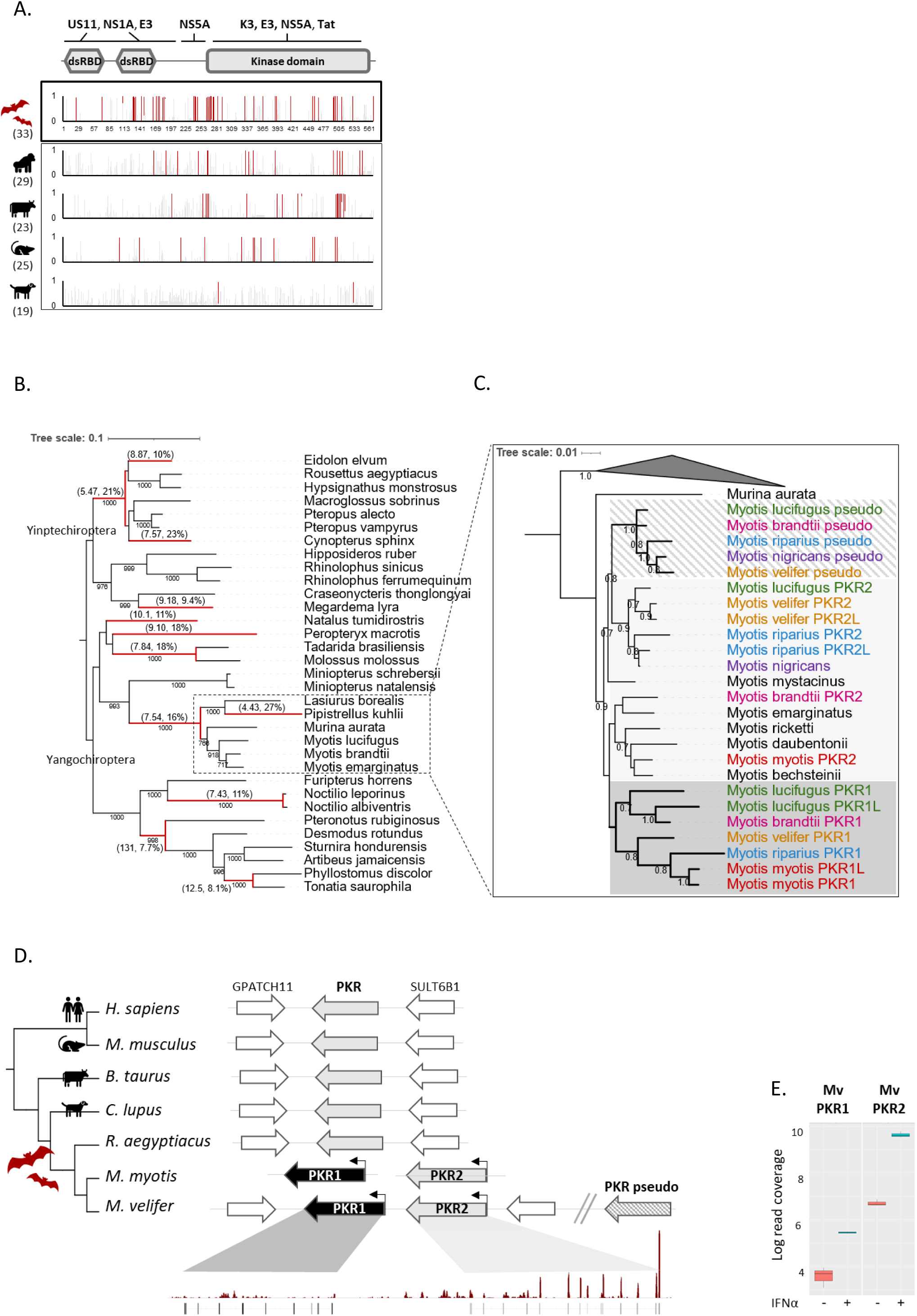
PKR has been the target of strong diversifying positive selection and original duplication in bats. **A.** Sites under positive selection in mammalian PKR. Graphic panels represent the posterior probabilities of positive selection (Bayes empirical Bayes, BEB) (y axis) in the M2 Codeml model (allowing for positive selection, ω >1) for each codon (x axis) in mammalian PKR. Red bars indicate the sites identified by both models, M2 and M8, with a BEB posterior probability greater than 0.95. Numbers in brackets are total species analyzed in each mammalian order (See Data availability section for alignments and trees). **B.** Maximum likelihood phylogenetic tree of bat PKR indicating the branches under significant positive selection (p-value <0.05, in red) assigned by aBSREL from the HYPHY package. The numbers in brackets indicate the estimated values of the ω at the branch. The scale bar indicates the number of substitutions per site . **C.** Maximum likelihood phylogeny of Myotis PKR paralogous transcripts, with *M. autata*, *E. fuscus*, *L. borealis* and *P. kuhlii* as outgroups (the three latter were collapsed to facilitate visualization). Colors indicate duplicated PKRs isolated from one species individual. Bootstrap values greater than 0.7 are shown. The scale bar represents The scale bar indicates the number of substitutions per site. **D.** Representation of the canonical locus of EIF2AK2/PKR gene in bats, primates, rodents, carnivores and artiodactyls. Plain colored arrows represent EIF2AK2 genes, the striped arrow shows the EIF2AK2 pseudogene, and white arrows are adjacent syntenic genes. The EIF2AK2 paralogous genes in *M. myotis* and *M. velifer* are located in tandem in the genome, while the pseudogene is located outside de canonical locus in the same chromosome. In *M. velifer*, the 5’UTR and first four exons were not found in the present genome. **E.** Expression pattern of PKR duplicates upon basal and IFNα treatment of *M. velifer* fibroblast cells. Boxplots represent number of reads in log10 scale for each condition, and each PKR copy. Analyses were restricted to the exons that are present in both PKR genes.

Other forms of genetic changes may be adaptive during evolutionary virus-host arms-races^28,29^. Notably, gene duplication and recombination are among the most important mechanisms underlying the diversification of the mammalian antiviral repertoire. We thus investigated how the gene encoding bat PKR, *EIF2AK2*, has evolved at the genomic level. Analyzing the publicly available genomes, we found distinct PKR-like sequences in the *Myotis* bats that suggested gene duplication of *EIF2AK2* specifically in this chiropteran lineage. However, because most of the publicly available PKR sequences from *Myotis* species are of low quality (i.e. highly fragmented, low coverage), and the PKR locus in the *M. myotis* genome^12^ is incomplete, we sampled seven new *Myotis* species (*M. bechsteinii, M. emarginatus, M. nigricans, M. riparius, M. myotis, M. mystacinus and M. velifer*; see Methods for details), and sequenced their complete PKR mRNA transcripts, as well as two genomic DNA fragments of the *EIF2AK2* locus. This allows identification of potential differences in intronic regions between the putative PKR duplicates, which would be evidence of authentic genomic duplication and not splicing variants. Combining our results from mRNA and gDNA data, we found that *EIF2AK2* has experienced repeated duplication events in a species-specific manner, leading to gene copy number variation (CNV) across *Myotis* species (Figure 1C, S2). In particular, we detected evidences of PKR duplicates in four *Myotis* species (*M. nigricans, M. riparius, M. myotis, M. velifer*), including a pseudogenized retrocopy of PKR that is specifically present in the New World clade of *Myotis* (Figure 1C, Figure S2-S6). *M. myotis* encodes two paralogous copies with intact open reading frames (referred to as PKR1 and PKR2), and one transcript variant (PKR1L) that may be a paralog or a splicing variant of PKR1 (Figure S3, S4, Figure 1C). In *M. velifer*, we isolated three distinct PKR sequences, two of which are paralogs (PKR1 and PKR2), while the third one is an isoform of PKR2 (Figure S3, S5). The same pattern was found in *M. riparius*, although the complete coding sequence of PKR was solely obtained for one copy, while the others were partial sequences. The other *Myotis* species (*M. bechsteinii*, *M. emarginatus*, *M. mystacinus)* had a single copy of PKR, although technical limits could have impaired the detection of PKR duplicates. The phylogenetic reconstruction of all PKR copies indicated that PKR has expanded before the diversification of the *Myotis* genus, 30 MYA ago, which was presumably followed by independent lineage-specific duplications (Figure 1C, D).

To characterize the genomic localization of *EIF2AK2* duplicates, we analyzed the genomic locus of *EIF2AK2* in *M. velifer* from an ongoing genomic sequencing project of the species (ongoing sequencing project by M. E. Lauterbur and D. Enard). We mapped two *EIF2AK2* copies in tandem in the *M. velifer* draft genome. However, while one copy had an integral structure spanning from the 5’UTR to the 3’UTR, the second gene lacked the 5’UTR and the first four exons, probably resulting from technical issues at the assembly step (Figure 1E, Figure S5). In addition, mRNA expression from RNAseq analyses of PKRs in *M. velifer* cells further showed that the two PKR copies are expressed in basal conditions and their expression is stimulated upon type-I IFNα treatment (Figure 1E).

### Genetic arms-races have shaped a specific bat PKR – poxviral K3 interface

Such extensive molecular and genomic changes in bat PKR could be the result of pathogenic virus-driven selective pressure. Specifically, because (i) we identified, in bat PKR, a hotspot of positive selection at residues known to interface with poxvirus antagonist K3 in primate PKR^19,24^, and (ii) poxviruses are currently circulating in bats^30–33^, we investigated the specificity of bat PKR-poxvirus K3 interface in heterologous virus-host assays^34^. On the virus side, we used a panel of K3 antagonists isolated from (i) Eptesipox virus (EPTV)^32^, which naturally infects the bat *Eptesicus fuscus*, (ii) the archetypal poxvirus vaccinia virus (VACV), and (iii) the well-known human pathogen variola virus (VARV). On the host side, we tested the bat PKR duplicated copies and seven orthologs from representative species of chiropteran divergence. We first used a surrogate yeast system in which the ability of PKR to drive translational shutoff in presence or absence of active antagonists can be directly assessed by measuring yeast growth rates ^19,24,35^. First, we found that the PKR paralogs and orthologs were all able to shut down protein synthesis in yeast, indicating that bat PKRs, including the duplicated PKR genes in *Myotis,* encode for functional proteins and retain their primary protein synthesis shutdown function (Figure 2A, Figure S7). Second, while the PKR paralogs had the same phenotype to K3 antagonism, the orthologous PKRs differed in their ability to escape poxviral K3s in a host species-specific manner (Figure 2A). Finally, we identified marked differences for PKR antagonism between the tested K3s, revealing virus-specific determinants of poxvirus K3 antagonism (Figure 2A). To test whether this was also the case in a mammalian cellular system, we used Hela PKR-KO cells in which we transiently co-expressed PKR +/- K3, as well as a luciferase expression-plasmid as a reporter system for cell translation. We obtained similar results (Figure 2C, D), thereby validating the reliability of our yeast system. In this assay, all PKRs showed comparably strong repression of luciferase expression, with the exception *M. velifer* PKR2, which showed somewhat weaker activity (Figure 2C).

**Figure 2.**
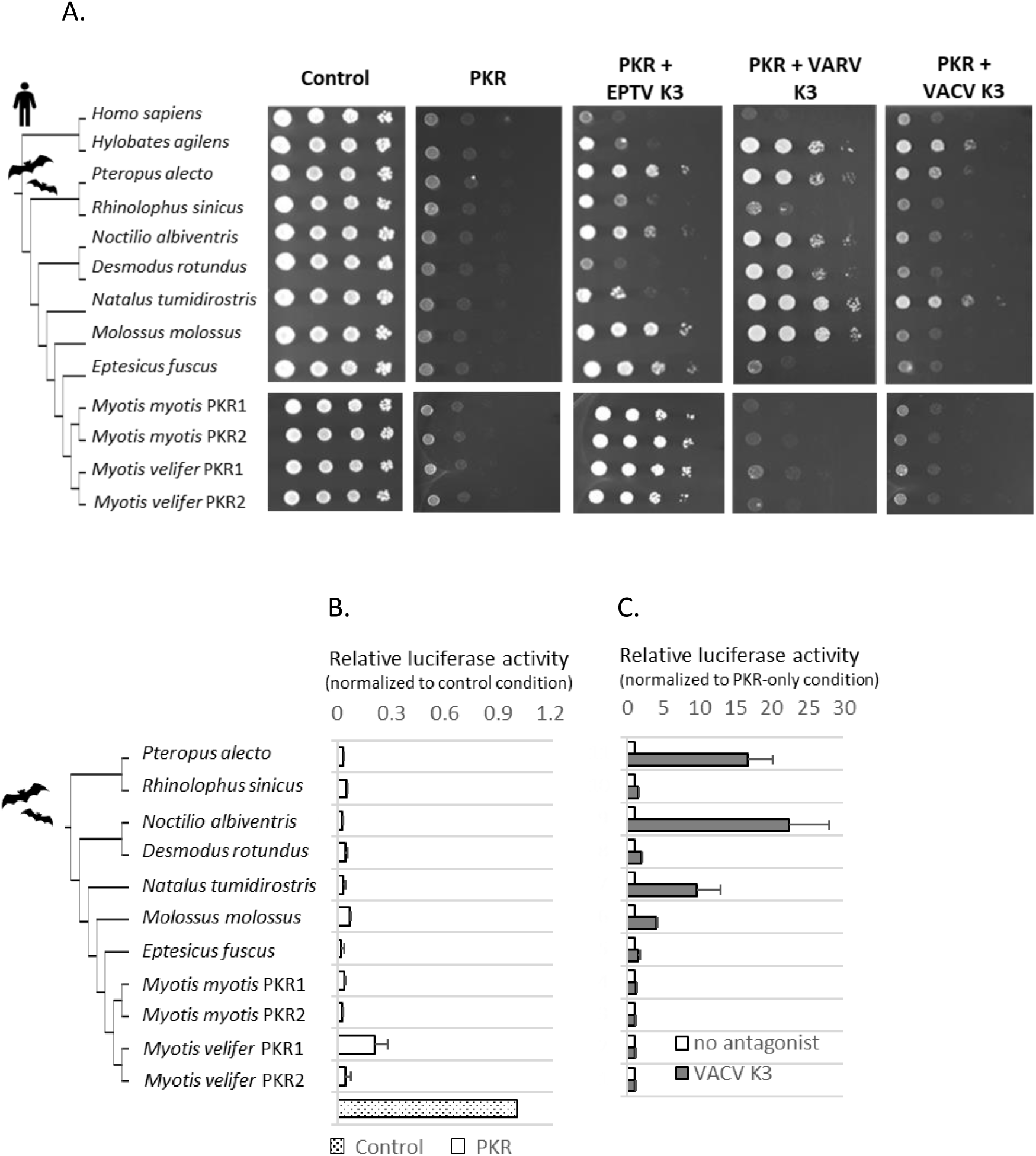
Species-specificity in bat PKR resistance and poxvirus K3 antagonism. **A.** Species-specific sensitivity of PKR to distinct poxvirus K3L proteins in yeast assays. Plasmids expressing PKR paralogous copies (PKR1 and PKR2) and orthologous variants under a galactose-inducible promoter were introduced into a wild type yeast strain or yeast strains expressing vaccinia, variola or eptesipox virus K3L. PKR variants from human and gibbon were used as positive control. Tenfold serial dilutions of transformants were spotted on plates containing either 2% glucose (control) or galactose. The Western-Blots for the expression of PKR and K3 proteins in yeasts are shown in Figure S6. **B.** Luciferase reporter assay showing that the PKR orthologs and paralogs inhibit the protein expression at comparable levels, except *M. velifer* PKR1 showing slight differences. Luciferase activity was normalized to the no-PKR condition in which cells were transfected with luciferase and empty vector **C**. Luciferase reporter assay confirming the differential sensitivity of PKR variants to vaccinia K3. Three independent experiments were conducted for bat PKR variants. Luciferase activity was normalized to the control condition in which cells were transfected with luciferase, PKR and empty vector (x axis). Error bars indicate Standard error of the mean, SEM.

To investigate the genetic determinants underlying these phenotypic differences, we used an evolutionary-guided approach on both the host and the virus sides. On the host side, because *D. rotundus* and *M. myotis* PKRs displayed opposite phenotypes to EPTV and VARV K3 antagonism, we generated a series of chimeras and mutants between these orthologs, and tested their capacity to escape EPTV and VARV K3 antagonism (Figure 2A, Figure S8). We showed that residues 475/476, located in the Helix αG in *D. rotundus,* drive species-specificity to variola K3 antagonism (Figure 2A). This determinant is similar to the previously reported residue 496 in human PKR-VARV K3 interface^24^. However, we further identified a yet undescribed within-protein epistatic^36^ interaction between the residues 475-476 and 332-344 in the kinase insert of *D. rotundus* PKR (Figure 2A) that represent specific determinants of susceptibility / resistance to EPTV K3. Swapping these sites significantly reduced K3-antagonist resistance of *D. rotundus* PKR, and conversely in *M. myotis* PKR, without impeding their expression and their basal translation shutdown function (Figure 2A, B). Importantly, these sites were among the fastest evolving codons in bat PKRs – with many substitutions and indels within the 332-344 amino acid stretch (Figure 2C) – significantly impacting the predicted 3D structure of bat PKR (Figure 2D). Therefore, accumulated mutations at these sites are adaptive in the context of bat-virus interactions and drive the host – poxvirus K3 specificity, supporting that ancient poxviruses that targeted these regions have been key drivers of inter-species bat PKR adaptation.

On the virus side, the protein alignment of orthopoxvirus K3 sequences revealed a unique structural C-terminal insertion in EPTV K3 (Figure 3E). To investigate whether this insertion functionally contributes to bat PKR antagonism, we tested the ability of an EPTV K3 mutant, which lacks the C-terminal insertion, to antagonize PKRs compared to the wild-type K3. We found that the truncated K3 had a significantly reduced anti-PKR activity, in a host-specific manner, without impacting K3 expression (Figure 3F). This shows that the C-terminal insertion in EPTV K3 is essential for bat PKR antagonism and accounts for species-specificity. Remarkably, combining these functional assays with a protein-protein docking model, which was performed with the HDOCK software^37^, we showed that the C-terminal insertion may be involved in PKR antagonism through direct contact with the residues 475-476 and 340, located in the Helix αG and the kinase insert, respectively (Figure 3G). In accordance with the functional assays, the protein complex between EPTV K3 and bat PKRs further depended on the bat PKR sequence and their 3D structure (Figure S9). Therefore, this C-terminal insertion may reflect counter-adaptation of EPTV K3 to maintain PKR antagonism.

**Figure 3.**
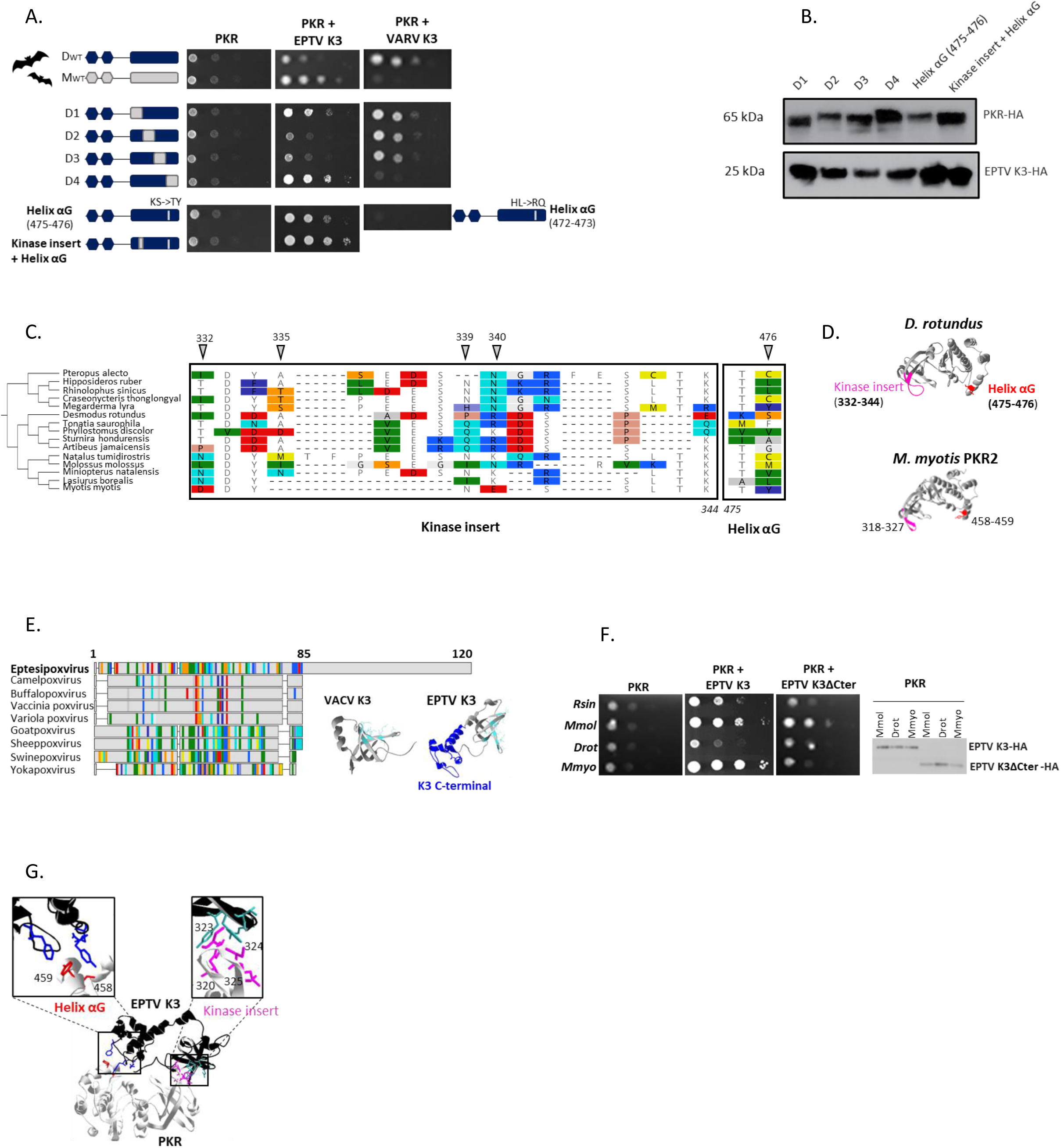
Evolutionary-guided functional approach reveals adaptive within-protein epistasis in bat PKR and a unique C-terminal extension in the bat poxvirus K3. **A.** Yeast spot assays of *D. rotundus* PKR mutants identifying the genetic determinants of PKR susceptibility to eptesipox virus and variola K3 antagonism. Mutants D1 to D4 are chimeric proteins between *D. rotundus* PKR and *M. myotis* PKR2, including the swap of amino acids 268-344, 345-380, 381-420, 421-530 in *D. rotundus* for D1, D2, D3 and D4, respectively. The Helix αG and kinase insert mutants were generated by site-directed mutagenesis of *D. rotundus* PKR, by swapping the corresponding residues from *M. myotis* PKR2. **B.** Representative western blot of PKR and K3 expression in yeast. **C.** Alignment of the residues underlying PKR-eptesipox virus K3 interface in the Helix αG and kinase insert of PKR. Left, species cladogram of the corresponding PKRs. Right, PKR protein alignment, colors indicate site variations between the sequences as compared to the consensus sequence with a threshold of 25% (Geneious, Biomatters; blue/red, hydrophilic/hydrophobic residues). The codon numbers are based on *D. rotundus* PKR sequence. The triangles indicate the residues under positive selection. **D.** 3D protein structure of *D. rotundus* and *M. myotis* PKR2 kinase domain, obtained by homology-modeling using I-TASSER and human PKR crystal structure (pdb 2a19). The residues identified by our yeast assays are colored in red in Helix αG and in magenta in the kinase insert. **E.** Multiple alignment and comparison of K3 protein sequence between divergent poxviruses. Colors indicate site variations between the sequences compared to the consensus sequence (threshold of 25%). Sequence numbering is based on eptesipox virus K3. In the right, 3D protein structure of vaccinia (pdb 1luz) and eptesipox virus K3 inferred by I-TASSER. The specific C-terminal insertion of eptesipox virus K3 is colored in dark blue. In light blue, residues involved in vaccinia K3-PKR binding^93^**F.** Yeast spot assays of bat PKRs* challenged with eptesipox virus WT and mutant K3ΔCter (Δ85-120 aa). Right, eptesipox virus WT K3 and K3ΔCter. **G.** Protein-protein complex structure between *M. myotis* PKR2 kinase domain and eptesipox virus K3, inferred by Hdock software. The kinase domain of PKR is represented in light grey and K3 in black. Residue coloring according to D (PKR) and E (K3) panels. *Rsin, *Rhinolophus sinicus;* Mmol, *Molossus molossus,* Drot, *Desmodus rotundus* ; Mmyo, *Myotis myotis*.

### The PKR paralogs functionally diverge in the ability to escape from poxvirus E3, cytomegalovirus TRS1 and influenza A virus NS1 antagonists

Because the PKR copies did not show phenotypic differences to poxvirus K3 antagonism, we tested whether they have evolved differences regarding their susceptibility to other poxvirus antagonists or to antagonists from others viral families that naturally infect bats or humans. In our experiments, we included (*i*) E3 antagonist from EPTV^32^ and VACV poxviruses, (i*i*) NS1 antagonist from human influenza A (IAV) H1N1 virus^38^, (*iii*) NS5A proteins from human hepatitis C virus (HCV) and from bat hepaciviruses infecting *Otomops martiensseni*^39^ (Omar) and *Peropteryx macrotis*^39^ (Pmac), and (*iv*) TRS1 from human cytomegalovirus^40^. Using the luciferase reporter assay, we showed that the *Myotis* bat PKR copies functionally differ in their ability to escape the viral antagonists NS1, TRS1 and E3 (Figure 4A-B, Figure S10). Although the immunoblot analysis showed that PKR2 was more expressed than PKR1 in this assay, this latter could efficiently escape from the viral antagonists. Importantly, in the case of VACV E3 antagonism, the strong intra-species differences between the PKR paralogs were of the order of magnitude of the inter-species PKR orthologs (Figure S11). Overall, this shows adaptive duplication of PKR in *Myotis* bats, which has broadened escape mechanisms to antagonism from very diverse viral families.

**Figure 4.**
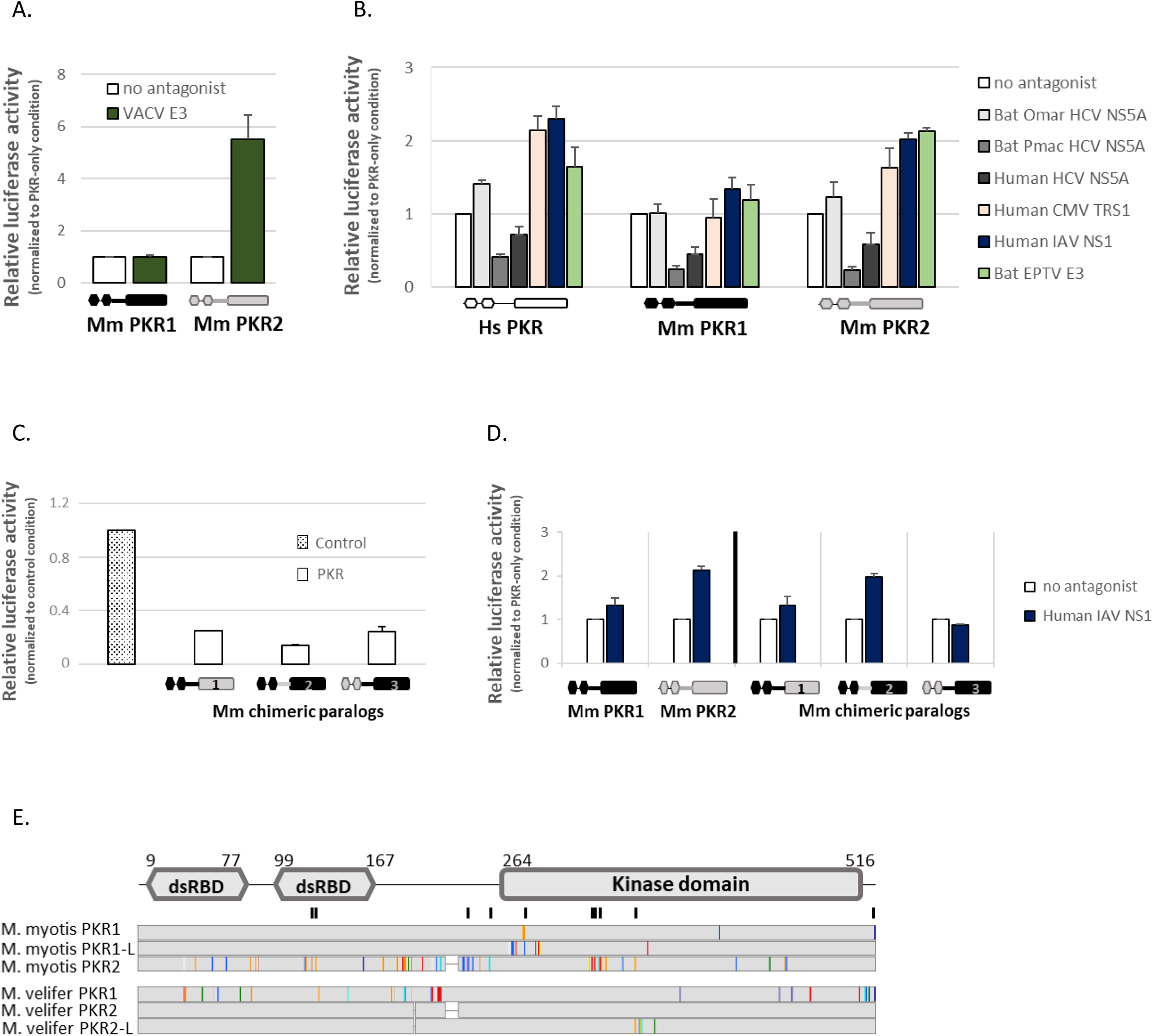
Functional divergence of the PKR paralogs in their ability to escape from poxvirus E3, cytomegalovirus TRS1 and influenza A NS1 antagonists. **A-B**. Relative luciferase activity in cells transfected with or without human PKR, *M. myotis* PKR1 or PKR2, in the presence or absence of putative viral antagonists: vaccinia virus E3 (**A**), and bat hepacivirus HCV NS5 (*O. martiensseni* and *P. macrotis* strains), human hepatitis C virus NS5 (JFH1), human Influenza A virus NS1, human cytomegalovirus TRS1, eptesipox virus E3 (**B**). The results shown are mean value of 3 and 5 independent experiments for panel A and B, respectively. Luciferase activity was normalized to the control condition in which cells were transfected with luciferase, PKR and empty vector. The error bars represent the SEM. **C**. Luciferase reporter assay showing similar translation inhibition by the *M. myotis* PKR paralog chimeras, which were generated by swapping the kinase domain (chimera 1), the linker region (chimera 2) or the dsRNA domain (chimera 3) of *M. myotis* PKR1 (black) with that of PKR2 (grey). Luciferase activity was normalized to the luciferase-only condition. The graph represents the mean of three independent replicates. **D**. Luciferase reporter assay showing the sensitivity of the PKR chimeras to human IAV NS1 antagonism (mean of three biological replicates). The error bars are the SEM. Luciferase activity was normalized to the condition without antagonist. **E.** Protein sequence alignment of *M. myotis* and *M. velifer* PKR duplicates and potential isoform. The black bars on top of alignment indicate the residues that differ between the paralogs and have evolved under significant positive selection. Colors indicate site variations as compared to the consensus within species with a threshold of 25%. The sequence numbering is based on *M. myotis* PKR1 sequence.

To decipher the underlying determinants of these functional differences, we engineered three chimeric PKR proteins, by swapping the dsRNA-binding domain, the linker region, or the kinase domain of the duplicated copies, and we tested their susceptibility to human influenza A NS1 antagonism. We showed that the linker region of the PKR paralogs drives the susceptibility or resistance to NS1, indicating that it is a key determinant for bat PKR antagonism by NS1 (Figure 4C, D). Remarkably, most of the genetic intra-species differences between the PKR copies are concentrated in the linker region (Figure 4E). Mapping the positively selected sites (inferred from the inter-species analyses) on the PKR paralogs, we found that several of these sites have undergone amino acid replacement in the PKR duplicates (Figure 4E), including in the linker region. Combined with our functional assays, these results indicate that (i) ancient viral pathogens from diverse RNA and DNA virus families may have contributed to the duplication and fixation of *Myotis* PKR paralogs, and (ii) the resulting evolutionary patterns in PKR paralogs account for distinct interactions with modern viral proteins.

### Bat PKR duplication leads to differential and potentially additive restriction of poxvirus and rhabdovirus infections

The fact that the PKR duplicated copies (i) inhibit cellular translation, (ii) are upregulated upon IFN stimulation, and (iii) are antagonized by diverse viral proteins suggest that both are potential antiviral restriction factors. To test this, we performed viral infection assays with a representative DNA virus and a representative RNA virus.

First, we created T-REx-293 PKR-KO cell lines expressing PKR1 or PKR2 from *M. myotis* or *M. velifer*, or *E. fuscus* PKR under doxycycline induction. For the infectivity assays with the DNA virus, we used a vaccinia poxvirus VC-R4^41^ lacking the viral K3 and E3 antagonists (VACVΔK3ΔE3) and expressing a virus-replication reporter EGFP. We found that the PKR paralogs significantly differed in their capacity to restrict VC-R4 replication. Whereas *M. myotis* and *M. velifer* PKR2 effectively inhibited VC-R4, as also seen for *E. fuscus* PKR, *M*. *myotis* and *M. velifer* PKR1 had only weak and no effect on the EGFP signals, respectively (Figure 5A-B). This was the case despite comparable expression of the PKR paralogs (Figure 5C). We further titrated VC-R4 replication in representative cell lines. T-REx-293 cells expressing *E. fuscus* PKR and *M. velifer* PKR2 were not included as they showed the same level of EGFP suppression as *M. myotis* PKR2. The titration supported the differences in EGFP signals in the microscopy images, with *M. myotis* PKR2 expression conferring a 1,000- fold reduction in VC-R4 titer, whereas only 3.6- and 1.2-fold titer reductions were observed for *M. myotis* PKR1 and *M. velifer* PKR1, respectively.

**Figure 5.**
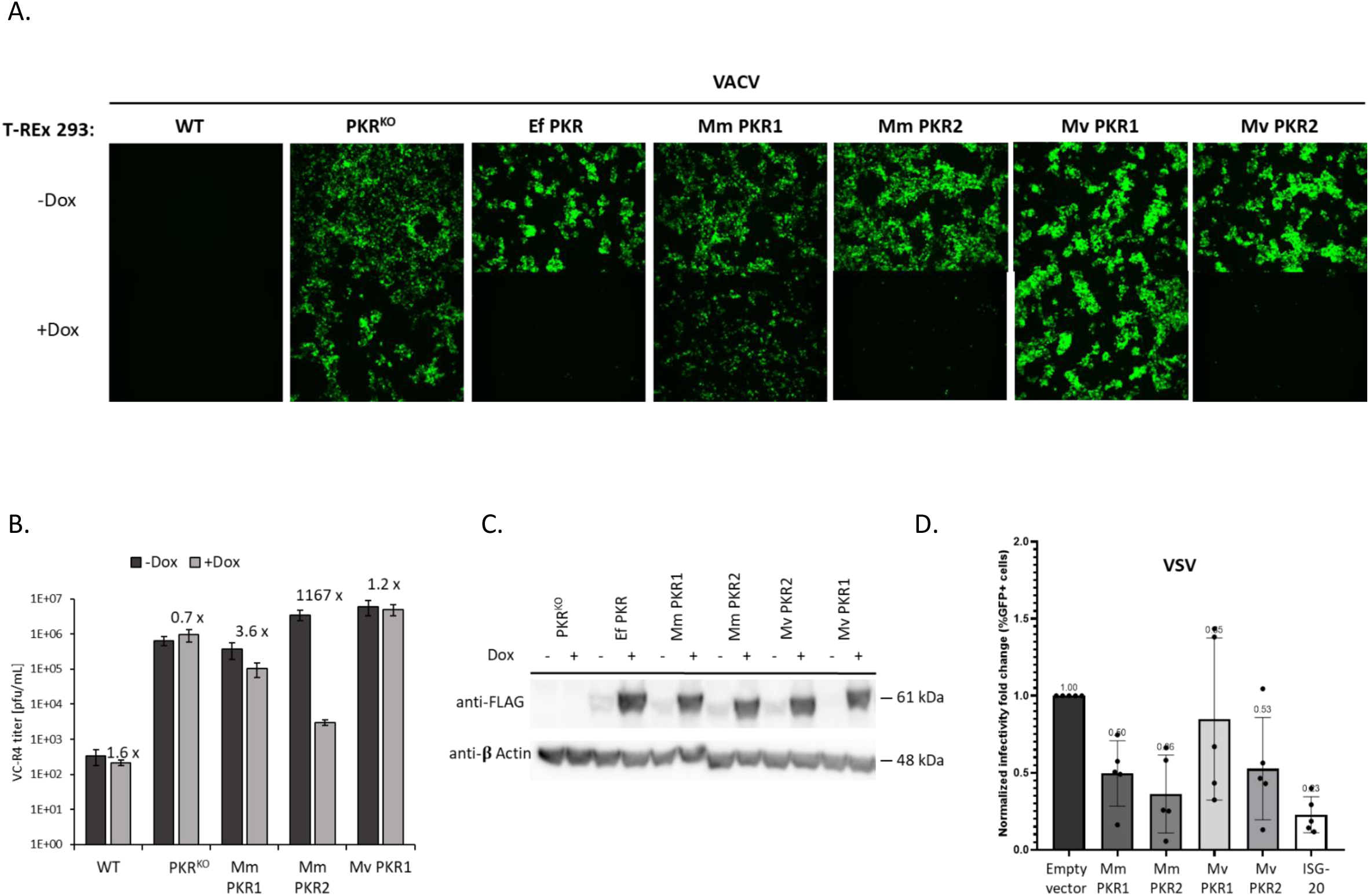
Bat PKR duplication allows for differential and potential additive antiviral restriction of poxvirus and rhabdovirus infections. **A.** T-REx-293 PKR-KO cells stably expressing *M. myotis* and *M. velifer* PKR 1 and 2, or *E. fuscus* PKR were infected with the VC-R4 (EGFP-VACVΔK3LΔE3L) at a MOI of 0.1, The EGFP expression were imaged 24h post-infection. **B.** Cells were infected as indicated above. Viruses in cell lysates were titered in RK13+E3L+K3L cells. Error bars represent the SD from two independent infections. Fold differences in virus titers obtained with -doxycycline and +doxycycline are shown. **C**. Expression of bat PKRs in the stably transfected T-REx-293 PKR-KO cells. Cell lysates were separated on 10% SDS-PAGE gels and analyzed by immunoblot analysis with anti-FLAG and anti-β-actin antibodies. **D.** Hela PKR-KO cells were transfected with or without *M. myotis* or *M. velifer* PKR1 or PKR2, or with ISG20 (as a positive control of viral restriction). Infection was performed 24h post-transfection with a VSV-GFP virus at MOI 3, and cells were fixed at 18hpi for flow cytometry analyses. Infectivity is measured as the ratio of the mean of % EGFP+ cells in each condition relative to the vector control condition. Values represent mean ± SD computed from 5 independent experiments.

Second, to determine whether this pattern was virus dependent, we further tested the antiviral function of the paralogs against an RNA virus, the vesicular stomatitis virus (VSV). We found that *M. myotis* and *M. velifer* PKR1s and PKR2s could all restrict VSV-GFP infection, although to varying extents (Figure 5D), indicating that *Myotis* bats have two VSV-restrictor PKR copies.

Therefore, PKR duplication may have contributed to the functional diversification and the potency of the *Myotis* antiviral repertoire, through distinct functional specialization of the PKR copies.

## Discussion

Combining in-depth phylogenetic and positive selection analyses with functional assays and experimental infections, we show how past genetic conflicts with pathogenic viruses have shaped chiropteran host antiviral immunity and susceptibility. In particular, we report extensive signatures of functional adaptation in PKR during bat evolution, with substantial molecular changes and genomic duplication, a novelty compared to other mammals. These adaptive changes now lead to species-specific interactions with contemporary viral pathogens and account for a specific, broad and potent antiviral response in bats.

In primate PKR, single substitutions at specific residues in the helix αG are key determinants for vaccinia and variola K3 antagonism^19,23^. In contrast, the genetic basis and specificity of bat PKR sensitivity to EPTV K3 relies on a within-protein epistatic interaction^36^ between two residues in the helix αG (475/476 in *D. rotundus*) and a stretch of amino acids in the kinase insert (332-344 in *D. rotundus*) of PKR. Although the role of this insert in PKR binding substrate was suggested in a previous study^42^, its functional implication in bat PKR–K3 interaction indicates that it contributes to substrate discrimination in bats. Under virus-host conflicts, the flexible and disordered feature of the kinase insert^43^ may have been a source of evolutionary plasticity allowing drastic changes in PKR while maintaining eIF2α binding. In line with this, repeated deletions/insertions were found in the kinase insert of several bat species without negative cost on basal protein shutdown function. Such hotspots of variability in unstructured loops are also found in other antiviral proteins and are prime targets of viral antagonism while being essential for antiviral activity^44,45^.

On the virus side, we showed that EPTV K3 evolved an adaptive Cter insertion that is essential for species-specific antagonism of bat PKR. This K3 Cter insertion was probably retained during EPTV evolution because of its increased PKR binding affinity through direct interaction with the kinase insert and the helix αG. Furthermore, because it increases PKR antagonism, the K3 insertion may not only drive the host range specificity, but may also directly or indirectly contribute to EPTV virulence in bat host species. Comparing the EPTV K3 sequence to all other available mammalian poxvirus K3s showed that this C-terminal insertion was specific to EPTV. Its 86 amino acid length suggests that it could derive from gene transfer, as frequently observed in poxviruses^46–48^. However, we failed to uncover the origin of this extension (i.e. no match in blat/blast searches), either from a parental host gene or from recombination of a viral sequence. One possible explanation is that the C-terminal sequence of EPTV K3 has substantially diverged from the parental one, such that their percentage of sequence similarity is negligible or non-existent. Further studies will be important to determine whether other bat poxviruses have evolved similar adaptive changes and decipher the functional implication in poxvirus pathogenicity and epidemiology.

Apart from poxviruses, other viral pathogens have certainly contributed to the diversification of PKR in bats. These mammals are highly diverse and have evolved with many viral pathogens over million years. Therefore, the evolution of their PKR may reflect the selective pressure of different ancient epidemics. Notably, the fastest evolving codons in bat PKR map with specific PKR-virus interfaces in primates, such as influenza virus NS1, cytomegalovirus TRS1, or hepacivirus NS5A ^18,40,49^, which homologs are encoded by bat-borne related viruses. Here, we found that ancient influenza- and cytomegalovirus-like viruses may have also been important drivers of PKR adaptation in bats, highlighting the diversity of viral selective pressures that have contribute to bat PKR evolution. In addition, the genetic differences between bat and other mammalian PKRs further suggest that specific bat-borne pathogens may be key actors and/or that related mammalian viruses may have evolved to antagonized different regions in bats.

Beyond substitutions or indels, we also found that gene duplication has diversified the bat antiviral repertoire in a lineage-specific manner. While all other studied mammals possess one single gene encoding PKR, several bat species from the *Myotis* genus express at least two functional, genetically divergent copies of PKR. Expansions of genes encoding antiviral proteins were previously discovered in bat species, including the APOBEC3, TRIM 22/5 and IFITM3 gene families, as well as the chimeric protein HERC5/6, or BST2^14,16,17,45,50^. Importantly, however, these duplications involve known multigene immune families, which are prone to gene expansion in many mammals, in contrast to the *EIF2AK2* (PKR) locus that is highly conserved in other mammals. Given the pleiotropic and central role of PKR in innate immunity, the duplication of PKR in *Myotis* bats reveals that major selective pressures have shaped bat evolution, leading to specific functional diversification in bat innate defense.

Prior to this study, independent PKR duplications were solely reported in amphibians and fishes^51–53^. In the latter, a PKR-like protein, containing a Z-DNA binding domain, was described as a cooperator of fish PKR antiviral activity^51,54,55^. In *Myotis* bats, the paralogs retained the typical structure of the mammalian PKR protein, with two dsRNA-binding domains linked to a kinase domain, but they genetically differ through multiple amino acid changes and indels. Because the evolutionary fate of gene duplication depends on the benefits and costs associated with the duplicated copies^56,57^, the fixation of PKR paralogs in *Myotis* genome suggests that they provide a functional selective advantage. Using two divergent RNA and DNA virus models (VSV and VACV, respectively) and various viral antagonists of PKR, we demonstrated that the PKR copies that could inhibit protein expression in a mammalian reporter assay, differed in their capacity to restrict virus replication and escape viral antagonism. Therefore, the PKR paralogs have retained the basal function of the parental copy (i.e. translation shutdown), but have evolved specific roles in the host antiviral response, as reported in other cases of gene duplication (e.g.^58,59^). One plausible explanation for this differential antiviral activity is that restriction potency depends on the virus. Alternatively, the PKR paralogs may have evolved to fill another functional niche. PKR is positioned upstream of several important factors, such as the immune transcriptional regulator IRF-1^60^, or the inflammatory transcription factors NF-kB^61,62^ and STAT1^63^. Losing one or some of these features could lower the overall antiviral response of one of the PKR paralogs. Finally, because PKR undergoes dimerization upon activation – which is essential for eIF2α phosphorylation^64^, it is possible that PKR1 and PKR2 heterodimerize to confer a new function. Although we only tested the PKR paralogs independently, this remains possible as we show that Myotis fibroblasts express the two genes. Thus, one could postulate a synergistic functional interaction between the paralogs upon viral infection, that could modulate their function.

Overall, this study brings important clues on the functional diversification of bat antiviral repertoire. It was suggested that immune tolerance rather than increased viral control plays a key role in bat immunity^12,65–68^. Here, the adaptive changes in bat PKRs increase the antiviral function and the viral evasion of PKR, which supports an adaptive enhancement for viral control in some species. This is in line with several studies reporting accelerated rate of evolution in bat restriction factors, indicating increased defense against virus infection^13,17,50,69^. Since each species has its own history of viral exposure, specific viral communities have certainly led to lineage-specific selection in bat’s antiviral immunity, highlighting the need to include multiple related species in comparative functional studies. Therefore, while dampening inflammatory response might be common to bats, strong episodic adaptations in antiviral factors, driven by ancient viral epidemics, may have shaped lineage-specific innate immune defenses in bats.

## Material and Methods

### Bat samples

Sampling was performed in France (*Miniopterus schreibersii, Myotis emarginatus, Myotis myotis, Myotis* and *Rhinolophus ferrumequinum*), French Guiana (*Desmodus rotundus*, *Myotis bechsteinii, Myotis riparius, Myotis nigricans, Peropteryx macrotis, Pteronotus rubiginosus, Tonatia saurophila, Natalus tumidirostris, Sturnira hondurensis, Molossus molossus, Noctilio albiventris and Furipterus horrens*) and Gabon (*Hipposideros cf. ruber, Rousettus aegyptiacus*). Authorizations were obtained from the Ministry of Ecology, Environment, and Sustainable development over the period 2015–2020 (approval no. C692660703 from the Departmental Direction of Population Protection (DDPP, Rhône, France). Our methods for animal capture and management were approved by the MNHN, the SFEPM and the DEAL-Guyane. African bat samples were approved by the Gabonese National Ethics Committee (Authorization N°PROT/0020/2013I/SG/CNE). Bat individuals were captured using harp traps at the entrance of caves or mist-nests hoisted on the forest floor and in the tree canopy. The individuals were then released after sampling. All samples, including wing membrane and blood pellet, were conserved at -80°C until RNA extraction.

In addition to the wild field samples, immortalized fibroblast cells from wing tissue of *Eptesicus fuscus* and embryonic fibroblast cell lines from *Myotis velifer* were generously provided by the Feschotte Lab (Cornell University)^70^. Cells were cultured in high glucose DMEM supplemented with 20% fetal bovine serum, 1% pen/strep and 1% sodium pyruvate.

### *De novo* sequencing of PKR (*EIF2AK2*) gene

Total genomic RNA was extracted from bat punches, fibroblast cells and blood samples using Macherey-Nagel Nucleospin RNA and RNA blood Kits, respectively, following the manufacturer’s protocol. Total RNA was reverse transcribed into complementary DNA (cDNA) with random primers and oligo(dT), using the SuperScript III One-Step RT-PCR reverse transcription kit (Thermo Fisher Scientific, Poland). Species identification was first confirmed through PCR amplification and sequencing of Cytochrome B gene (Cytb), using the primers CytB-F and CytB-L/R^71^ and the PCR protocol in Table S2. PKR mRNA was then amplified from each species using 30ng of cDNA and different sets of primers (Table S3) that were specifically designed using an alignment of publicly available PKR sequences. The PCR mix and conditions are presented in Table S2. PCR products with multiple bands were excised and purified from gel using the Nucleospin Gel and PCR Clean-up kit from Macherey-Nagel, or cloned using the NEB**^®^** PCR cloning kit (New Englands BioLabs) to obtain haplotype resolution. Sanger sequencing of PKR was performed by a commercial company (Genewiz, Azenta Life Sciences, Germany).

### Collection of PKR orthologous sequences

To complete our dataset, orthologous coding sequences of bat PKR were retrieved from Genbank by BLASTn searches of the “Nucleotide”, “Refseq Genome”, “Transcriptome Shotgun Assembly” and “Whole-Genome Shotgun Contigs” databases, using the Little Brown bat (*Myotis lucifugus*) Refseq coding sequence as query. In the case of unassembled bat genomes, PKR coding sequence was predicted from the genome contigs using Augustus^72^ and GeneWise^73^ with the Little Brown bat Refseq protein as reference. In total, 19 bat PKR sequences were retrieved from public databases.

PKR coding sequences from primates (n=29), rodents (n=25), artiodactyls (n=23) and carnivores (n=19) were obtained by tBLASTn searches of the “Nucleotide” database from Genbank using human (*Homo sapiens*), mouse (*Mus musculus*), cow (*Bos taurus*) and dog (*Canis lupus familiaris*) PKR protein sequence as queries, respectively.

### Phylogenetic and positive selection analysis of PKR orthologous sequences

PKR orthologous codon sequences were aligned for each mammalian group separately using the program PRANK^74^, and the alignments were manually curated. We then built a phylogenetic tree, using the maximum likelihood method implemented in PhyML program^75^. Selection of the best substitution model was performed with the Smart Model Selection (SMS)^76^ program in PhyML and was always: GTR+G+I. Node statistical support was computed through 1,000 bootstrap replicates. The detection of recombination events was assessed with GARD^77^.

For positive selection analyses, models that disallow positive selection (models M1 and M7) were compared to those allowing for positive selection (M2 and M8) using the PAML Codeml package^27^, with the following parameters: codon frequencies F61 and F3×4 and starting omega ω (*dN*/*dS* ratio) of 0.4. Comparison of each pair of models (M1 vs M2, and M7 vs M8) was then achieved with likelihood ratio tests. Bayes Empirical Bayes (BEB) of the *dN*/*dS* >1 class in M2 or M8 models was used to assess positive selection at the codon level, with a posterior probability ≥ 0.95 as significance threshold. The Fast-Unbiased Bayesian Approximation (FUBAR)^78^ and the Mixed Effects Model of Evolution (MEME)^79^, both implemented in the HYPHY package, were also run to identify codons under significant positive selection. To ensure higher specificity, we considered that codons were under significant positive selection if they were identified by at least two methods. Moreover, to test if the PKR domains (i.e. the dsRBD, the linker region and the Kinase Domain) have similarly been targets of positive selection, each domain was separately analyzed using the models M1, M2, M7 and M8 from the PAML package.

Finally, we determined if and how PKR experienced episodic selection during bat and mammalian evolution, using the branch-specific analysis aBSREL^80,81^, implemented in the HYPHY package. This program allows testing the significance of positive selection and quantifying the *dN*/*dS* ratio for each branch independently. Sequences from perissodactyls (n=3) and proboscidean (n=1) were also analysed. Tree visualization and annotation were performed with iTOL webserver (https://itol.embl.de/).

### Genomic and (phylo)genetic characterization of PKR paralogs in Myotis

Molecular identification of EIF2AK2 duplication was carried out in tissues from *Myotis* species, including *M. myotis*, *M. velifer*, *M. riparius*, *M. nigricans, M. mystacinus, M*. *emarginatus*, and *M. bechsteinii*. Total RNA and Genomic DNA were extracted using the Macherey-Nagel Nucleospin RNA tissue and gDNA kits, respectively, following the manufacturer’s instructions. Two complementary strategies were then used. First, PKR coding sequence was amplified from cDNA using the PKR “universal” primers designed in this study (Table S3). Second, from gDNA, we PCR amplified the genomic regions containing exons 1 to 3 (E1-E3), and exons 4 to 6 (E4-E6), of EIF2AK2 to identify potential differences in intronic regions between the putative PKR duplicates. Following PCR amplification, all PCR products from cDNA and gDNA were cloned into the pMiniT 2.0 Vector using the NEB cloning kit (New Englands BioLabs) and sequenced to ensure sequencing of a single DNA molecule.

Phylogenetic reconstruction of the PKR paralogs followed the previously-described phylogenetic analysis method.

We combined different methods to map and predict the EIF2AK2 locus in the *M. velif*er genome. First, we performed a BLASTN search (cut off 10^-05^) with PKR cDNA sequences from related *Myotis* species to identify the canonical locus and localization of EIF2AK2 gene copies. Second, we aligned sequences of proteins and RNA transcripts of PKR from related bat species on the *M. velifer* genome using the Fast Statistical Alignment (FSA) software^82^. Third, we integrated RNA sequencing (RNAseq) data (see methods below), by mapping the RNA-seq reads using HISAT2 (v.2.0.0)^83^ . Finally, we *de novo* predicted the gene structure of each PKR copy using Augustus^72^ in single-genome mode with the human gene model. The final figure was generated with the R library, Gviz.

### Interferon (IFN) cell treatment and transcriptomic analyses

*M. velifer* cells were seeded in 6-well plates. Forty-eight hours later, they were treated or not with 1,000U/ml of type-I universal IFN (pbl Assay Science). Six hours post-treatment, cells were collected and total RNA was extracted using Macherey-Nagel Nucleospin RNA. Six sample replicates (three without interferon treatment and three with) were then sent for library preparation and sequencing with Illumina NextSeq500, 150-paired-end, to the IGFL (*Institut de génomique fonctionnelle* de Lyon) sequencing platform.

We processed the RNA-seq data with the non-annotated draft *M. velifer* genome. The quality of the raw data was checked with FastQC and a Q20 threshold, and adapters were removed using Cutadapt 4.0^84^. The quality-controlled reads were then aligned to the *M. velifer* genome using HISAT2^83^. As a complement, we de novo predicted the gene from *M. velifer* genome using Augustus, with the human model as reference. We counted the number of reads that mapped to each gene (predicted by Augustus), in both basal and IFN conditions, with FeatureCounts (from the R package, RsubRead), and performed a differential analysis using DESeq2^85^. From this analysis, we obtained a list of genes with a significantly different number of reads between the two conditions. Because the genomic analysis of *M. velifer* showed a possible sharing of 5’ exons between the paralogs, we retained the number of reads from exons that were duplicated and specific to each paralog to assess the expression pattern of the PKR copies. The final figure representing the number of reads per paralog per condition was drawn via the R package, ggplot.

### Protein structure prediction and docking models

The 3D protein structures of *D. rotundus* and *M. myotis* PKR kinase domain, as well as the eptesipox virus K3 protein, were predicted using the Iterative-Threading ASSEmbly Refinement (I-TASSER) server^86–88^. The best model was carefully chosen based on the C-Score which assesses the quality of the models. The inferred protein structure of PKR was visualized and designed with Swiss PDB viewer software^89^.

Computational docking of bat PKR and eptesipox virus K3 was performed to predict the complex structure between both proteins, using HDOCK webserver^37^. This software uses a fast Fourier transform (FFT)-based search strategy to model different potential binding means between the proteins, then each binding mode is evaluated using the scoring function ITScorePP. The 3D structure models of *M. myotis, D. rotundus* and *M. molossus* PKR kinase domains, as well as eptesipox virus K3, were obtained with I-TASSER. We kept the default parameters for computation, including a grid spacing set to 1.2 Å and the angle interval set to 15∘A. We retained the first top three models and combined the docking results with our functional assays for final model selection.

### Plasmids

#### Expression in yeast cells

Bat eptesipox virus^32^ (Washington strain) K3L and E3L sequences were synthetized (Genewiz) with an integrated C-terminal HA-epitope tag, and cloned into the yeast LEU2 integrating plasmid pSB305 which contains a galactose promoter, using the *Xho*I and *Not*I restriction sites. Bat PKR cDNAs from divergent chiropteran families (*P. alecto, R. sinicus, D. rotundus, N. tumidirostris, M. molossus, N. albiventris, M. myotis, E. fuscus*) were cloned into the yeast pGAL expression plasmid, pSB819 (URA), using the *Xho*I and *Not*I restriction sites. The human and gibbon PKR expression vectors were previously described^23^. *D. rotundus* × *M. myotis* PKR chimeras were synthetized and sub-cloned into pSB819. PKR site-specific mutants and epteK3Δ227-508 were generated by PCR mutagenesis using the QuickChange Lightning mutagenesis kit (Agilent) and primers holding the desired mutations/deletions, following the manufacturer’s protocol. PKR mutants and chimeras were amino-terminal HA-tagged by PCR, using integrated HA-tag PKR primers (Table S3).

#### Expression in human cells

Bat PKR orthologs and paralogs were sub-cloned from pSB819 into the expression vector pSG5, by means of *KpnI* and *XhoI* sites introduced into PKR primers. Cloning of human PKR was previously described^19^. Full length NS5A proteins from human Hepatitis C Virus (JFH1), bat hepacivirus from *O. martiensseni* (NC_031947.1^39^) and bat hepacivirus L from *P. macrotis* (NC_031916.1^39^) were synthetized and cloned into the expression plasmid pCDNA3.1+ N-Terminal HA-tag using *BamhI* and *NotI* restriction sites. The eptesipox virus E3 and vaccinia E3 genes were cloned into the expression vector pSG5. The human Influenza A virus A/England/195/2009(H1N1) NS1 expressing-plasmid (pCAGGS V5-tag)^38^, as well as the human cytomegalovirus TRS1 plasmid (pCDNA3.1 HA-tag)^40^, were kindly provided by Wendy Barclay and Adam Geballe, respectively.

### Yeast strains and growth assays

To determine whether bat PKR variants differed in their ability to escape poxviral antagonism, we used a heterologous yeast growth assay^35^. This method relies on the recognition and phosphorylation of yeast eIF2α by PKR, which leads to yeast growth arrest. However, co-expression with poxvirus K3 or E3 that are able to antagonize PKR leads to growth rescue. Yeast growth assays were performed in two steps.

First, yeast strain H2557 was modified for stable expression of bat poxvirus K3 and E3 proteins following standard yeast transformation protocol^90^. Eptesipox virus K3 (eptK3), or eptesipox virus E3 (eptE3) was integrated into H2257 at the LEU2 locus under the gal promoter, using subcloned pSB305 plasmids linearized with EcoRV. The resulted strains H2557-eptK3, H2557-eptE3, and H2557-pteE3 were confirmed through PCR amplification and sequencing of K3 and E3, using the universal primers M13 F and M13R. Yeast strains expressing vaccinia and variola HA-K3, as well as the wild-type control (HM3, with integrated empty vector) were previously described^24^.

Second, the modified yeast strains were transformed with 100ng of PKR expression plasmids pSB19. For each transformation, four colonies were selected and streaked on S-leu-ura medium (yeast minimal complete medium with amino acids minus uracil and leucine) containing 2% glucose (SD) or galactose (SGal), and grown at 30°C for 3 days.

Representative transformants colonies were then grown to saturation in SD-leu-ura medium and plated in dilution series (D600 3.0, 0.3, 0.03, 0.003) on SD and SGal-leu-ura medium for 3 days. All yeast assays were conducted in biological triplicate experiments.

### Cell lines

HeLa PKR-knockout cells (kindly provided by Adam Geballe^40^) were maintained in Dulbecco’s Modified Eagle’s Medium (DMEM) supplemented with 5% fetal bovine serum and 1 μg/ml puromycin (Sigma). RK13+E3L+K3L cells (rabbit)^91^ were maintained in DMEM supplemented with 5% fetal bovine serum, 100 IU/mL penicillin/streptomycin, 500 μg/mL geneticin and 300 μg/mL zeocin (Gibco). Wildtype (Invitrogen) and PKR-KO T-REx-293 cells (Rothenburg lab, unpublished) were grown in DMEM supplemented with 10% fetal bovine serum, 100 IU/mL penicillin/streptomycin, 100 μg/ml zeocin and 15 μg/ml blasticidin (Gibco). The T-REx-293 cells stably transfected by bat PKRs were under constant selection ofwith 15 μg/ml blasticidin and 50 μg/ml Hygromycin (Invitrogen).

### Luciferase Reporter Assays

Luciferase assays were carried-out following the protocol described in^21^. Briefly, 50,000 Hela PKR-KO cells were seeded per well in 24-well plates, and transfected 16h post-seeding with 350 ng of PKR expression vector or empty control, 350 ng of viral antagonist expression plasmid (NS1, NS5A, EPTV E3 or TRS1) or empty control, and 5 ng of FFLuc firefly luciferase reporter plasmid, using Trans-IT-LT1 (Muris Bio) following the manufacturer’s protocol. Cells were lysed 48h post-transfection by means of the reporter lysis 5X buffer (Promega), then the luciferase substrate (Promega) was added following the manufacturer’s recommendations. Luciferase reporter quantitation was carried out with a LUMIstar omega microplate reader optima (BMG Labtech). All luciferase assays were conducted in triplicate in at least five independent experiments. For the luciferase assays with VACV K3 and E3 antagonists, 50,000 Hela PKR-KO cells per well were transfected 24h post-seeding with 200 ng of PKR expression vector, 200 ng VACV E3 expression plasmids (VACV K3 and E3), 50 ng of pGL3 firefly luciferase expression vector (Promega) using GenJet (Signagen) at a DNA to GenJet ratio of 1:2 following the manufacturer’s protocol. Cells were lysed 48h post-transfection with mammalian lysis buffer (GE Healthcare), then the luciferase substrate (Promega) was added following the manufacturer’s recommendations. Luciferase reporter quantitation was carried out with a Glomax luminometer (Promega). All luciferase assays were conducted in triplicate in at least three independent experiments.

### Generation of doxycycline-inducible bat PKR-expressing 293 cells

Bat PKRs (*E. fuscus*, the two *M. myotis* paralogs and the two *M. velifer* paralogs) were cloned into the pcDNA5/FRT/TO expression vector with two C-terminal FLAG tag sequences. T-REx-293 PKR-KO cells were stably transfected with each bat PKR plasmid by GenJet (Signagen) according to the manufacturer’s instructions and polyclonal pools of the stably transfected cells were selected by their resistance to hygromycin.

### Poxvirus infection

Generation of VC-R4, a derivative of VACV Copenhagen strain, was described^41^. 500,000 of T-REx bat PKR expressing cells were seeded per well in 12 wells plates and induced with 1 μg/mL doxycycline for 24h. 48h post-seeding, each well was infected by VC-R4 at MOI of 0.1. Fluorescent pictures were taken with an inverted fluorescent microscope (Leica) at the indicated time post infection. For the virus replication assay, cells and supernatants were collected 24h post-infection and subjected to three rounds of freezing at -80 °C and thawing at 37 °C. Lysates were sonicated for 15s, 50% amplitude (Qsonica Q500). Viruses were titered by 10-fold serial dilutions on confluent RK13+E3L+K3L cells in 12-well plates. One hour after infecting RK13+E3L+K3L cells with the dilutions, the medium was replaced with DMEM containing 1% carboxymethylcellulose (CMC). After 48 hours, cells were stained with 0.1% crystal violet and counted for plaques. Infections and viral titer were performed in duplicate.

### VSV infection

200,000 Hela PKR-KO cells were seeded per well in 12-well plates and transfected with either empty pSG5 plasmid, *M. myotis* PKRs or *M. velifer* PKRs, using Trans-IT-LT1. ISG20 encoding plasmid^92^ was used as a positive control, because of its established antiviral activity against VSV. Infection was performed 24h post-transfection, with a VSV-GFP virus at MOI3^92^, and cells were fixed at 18hpi with paraformaldehyde (PFA) 4%. Single cell analysis was performed using BD FACSCanto™ II Flow Cytometer to quantify VSV infectivity as the percentage of GFP+ cells. Fold-change results are normalized to the empty (no PKR or ISG20) condition from five independent experiments.

### Western blots

To examine the yeast expression of bat PKR and poxvirus K3 proteins, yeast transformants were grown overnight in 2% glucose S-leu-ura medium, followed by induction with 2% galactose for 15 h. Cell lysates were treated with 0.1M of Sodium Hydroxide (NaOH) for 5 min, then lysed in 2X SDS-PAGE buffer supplemented with protease inhibitor cocktail (Roche) and 355 mM β-mercaptoethanol (Sigma) at 95°C for 5 min. PKR was then precipitated at 65°C for 45 min, frozen overnight, and re-precipitated. Proteins were resolved by 12% Mini-PROTEAN GTX polyacrylamide gel (Bio-rad), then transferred to PVDF membranes. Proteins were probed with rabbit anti-HA (1:1,000 Sigma-Aldrich, H3663) or anti-β-actin as loading control (1:10,000 Sigma-Aldrich, A5441) primary antibody, then with goat anti-rabbit secondary antibody. Blots were visualized using the Image Lab Touch Software (version 2.0.0.27, Chemidoc Imaging System from Bio-Rad) or film.

In HeLa-KO cells, protein expression was assayed with 400,000 cell per well in six-well plates and transfected with 1.4 μg of the indicated PKR and viral antagonist expressing-plasmids. Cells were lysed after 48h with 1% sodium dodecyl sulfate (SDS) in PBS (VWR), then proteins were separated on 4-12% Mini-PROTEAN GTX polyacrylamide gel (Bio-rad) and transferred to nitrocellulose membranes. Proteins were resolved and visualized as described above.

To detect the expression of bat PKRs in the stably transfected T-REx 293 cells, 600,000 cells were seeded per well in six-well plates and induced by 1 ug/mL of doxycycline 24h post seeding. Cells were lysed 24h post induction with 1% SDS, then proteins were separated on 10% TGX Fastcast Acrylamide gel (Bio-rad) and transferred to PVDF membranes. Proteins were probed with mouse anti-FLAG (1:5,000 Sigma-Aldrich, F3165) or anti-β-actin (1:5,000 Sigma-Aldrich, A1978) primary antibody, then with donkey anti-mouse (1:10,000 Fisher Scientific, 715-035-150) secondary antibody. Images were taken using the iBright Imaging System (Invitrogen).

### Statistical analyses

Expression data were analyzed using Student t-tests and ANOVA, followed by Tukey’s post hoc test for pairwise comparisons, using R. For each pairwise comparison, we ensured that normality and homoscedasticity assumptions were met for the residuals using, respectively, Shapiro-Wilk and Levene tests. For each of these tests, the *p-value* was considered significant when inferior to 0.05. Error bars in graphics are Standard Error of the mean (SEM) .

## Data availability

All *de novo* PKR sequences were deposited in GenBank under Accession Numbers [waiting for accession numbers]. Dataset for Figure 1 with alignments and phylogenetic trees are publicly-available at https://figshare.com/projects/Datasets_for_Jacquet_et_al_2022/142388.

## Acknowledgments

We are particularly grateful to the Poitou-Charentes association, as well as the volunteers and field workers who have helped us during the field sessions: V. Alt, M. Bely, G. Chagneau, M. Dorfiac, S. Dufour, C. Gizardin, G. Leblanc, M. Leuchtmann, E. Loufti, A. Le Guen, and L. Trebucq. We thank B. Larsen, M. Bucci, and R. Ledgister for help with Myotis velifer sample collection, Pima County, AZ, USA for sampling permission, and Dovetail Genomics for sequencing. For their advices and the RNAseq analysis that we used for the PKR transcripts, we thank Marie Sémon, Marie Cariou, Carine Rey, Corentin Dechaud, Romain Bulteau, the students from the Master UE NGS, and the Master Biosciences, École Normale Supérieure de Lyon, Université Claude Bernard Lyon 1, Université de Lyon, 69342 Lyon Cedex 07, France, as well as Benjamin Gillet and Sandrine Hughes from the IGFL platform for the library preparation and the sequencing. We thank Cédric Feschotte and Rachel Cosby (Cornell University, NY) for generously sharing *Myotis velifer* and *Eptesicus fuscus* cell lines. We also thank Adam Geballe (Fred Hutchinson Cancer Center, WA) and Wendy Barclay (Imperial College London, UK) for sharing reagents (the references are given in the manuscript). We acknowledge the contribution of the SFR Biosciences (UAR3444/CNRS, US8/Inserm, ENS de Lyon, UCBL) ANIRA cytometry platform, especially Véronique Barateau for her help. We also thank all the contributors of the LBBE bioinformatic server, of the publicly available bioinformatic programs, and of the publicly available genomic sequences.

## Author contributions

Study conceptualization: SJ, LE, DP. Methodological design: SJ, SR, NE, DP, LE. Field sampling: J-BP, OFC, BN, JD. Phylogenetic and genomic analyses: SJ. RNAseq processing and analysis: SJ, AEF, CD, LE. Technical support: CD. Genome sampling and sequencing: MEL, DE. Yeast assays: SJ, MC, CV, CC. Luciferase assays: SJ, CZ, CP. Infectivity assays: CZ, CDLM, GB. Study coordination and supervision: LE, DP. Further supervision: SJ, SR, NCE. Funding: SJ, LE, DE, DP, AC, SR, NCE. Writing – original draft: SJ. Writing – first editing: LE, DP. Writing – Review and editing: all the authors.

## SUPPLEMENTARY

### Supplementary Tables

**Table S1.** Results of the positive selection analyses comparing models that disallow positive selection (M1 and M7) to models allowing positive selection (M2 and M8). ^a^ p-values generated from maximum likelihood ratio tests indicate whether the model that allows for positive selection (models M2 and M8) better fits the data than the nearly neutral one (M1 and M7). ^b^ Percentage of codons evolving under positive selection (dN/dS ratio > 1 over the alignment). -, not significant. ^c^Average dN/dS ratio associated with the classes K3 and K11, in the Codeml models M2 and M8, respectively, which allow positive selection.

|  | Codeml M1 vs M2 |  |  | Codeml M7 vs M8 |  |  |
| --- | --- | --- | --- | --- | --- | --- |
|  | p-value<sup>a</sup> | % of PSS<sup>b</sup> | M2 ω<sup>c</sup> | p-value<sup>a</sup> | % of PSS<sup>b</sup> | M2 ω<sup>c</sup> |
| **Chrioptera** |  |  |  |  |  |  |
| Whole gene | 4,4E-83 | 17 | 3.08 | 7,7E-86 | 20 | 2.76 |
| dsRNA domains (9-166) | 2,2E-22 | 16 | 3.11 | 3,9E-25 | 20 | 2.73 |
| Linker (167-265) | 9,8E-27 | 23 | 3.60 | 2,5E-27 | 23 | 3.57 |
| Kinase domain (266-536) | 2,7E-44 | 16 | 3.26 | 6,0E-46 | 19 | 2.92 |
| **Carnivores** | 4,8E-30 | 25 | 1.78 | 2,7E-31 | 28 | 1.72 |
| Cetoartiodactyls | 4,2E-04 | 13 | 4.30 | 2,5E-05 | 13 | 4.31 |
| Primates | 4,8E-30 | 16 | 3.29 | 2,7E-31 | 19 | 3.09 |
| Rodents | 1,4E-24 | 7 | 2.90 | 1,2E-25 | 11 | 2.25 |

**Table S2.** Results from site-specific positive selection analyses, with a posterior probability (PP) of BEB > 0.95 for the models M2 and M8 from Codeml, > 0.9 for FUBAR, and a p-value of 0.05 for MEME. Codons in bold were assigned by at least two methods. Codon numbering is based on *D. rotundus*.

| M2 | M8 | FUBAR | MEME | FEL |
| --- | --- | --- | --- | --- |
| 24, 26, 28, 72, 86, 97, **108**, 122, 123, 124, 125, 127, **138**, 143, 159, 165, **169**, **171**, 177, **179**, 181, **228**, 230, 231, 235, 251, 253, 256, **258**, **260**, 263, 264, 271, 279, 316, 332, **340**, **364**, **367**, 393, **431**, 433, 437, 444, 469, 470, **473**, **476**, **484**, 505, 536 | 7, 24, 26, 28, 72, 86, 97, 107, **108**, **114**, 120, 122, 123, 124, 125, 127, **138**, 143, 159, 165, **169**, **171**, 177, **179**, 181, 182, 221, **228**, 230, 231, 235, 250, 251, 253, 256, **258**, **260**, 263, 264, 267, 271, 279, 316, 320, **330**, 332, **335**, **339**, **340**, 360, **364**, **367**, 370, 373, 384, 393, **431**, 433, 437, 444, 469, 470, **473**, **476**, **484**, 505, 536 | 7, **81**, **108**, 135, **138**, **169**, **171**, **179**, **228**, **258**, **260**, **330**, **335**, **339**, **340**, **364**, **367**, **473**, **476**, **480**, **484**, 500 | 22, 74, 75, **81**, 82, 107, **108**, 111, **114**, 119, 128, **135**, 151, **169**, **171**, **179**, 200, **228**, 242, **258**, **260**, 265, 270, 289, 301, 318, 320, 327, **330**, **335**, **339**, **340**, 370, 380, 406, 427, **431**, 445, **476**, 477, **480**, **484**, 500, 508, 511 | **81**, 84, **108**, 111, **114**, 119, **135**, 151, **169**, **171**, **179**, 216, **228**, 232, 254, **258**, **260**, 328, **335**, **339**, **340**, 367, 384, 408, **476**, 477, **480**, **484**, 490, 508, 513 |

**Table S3.**
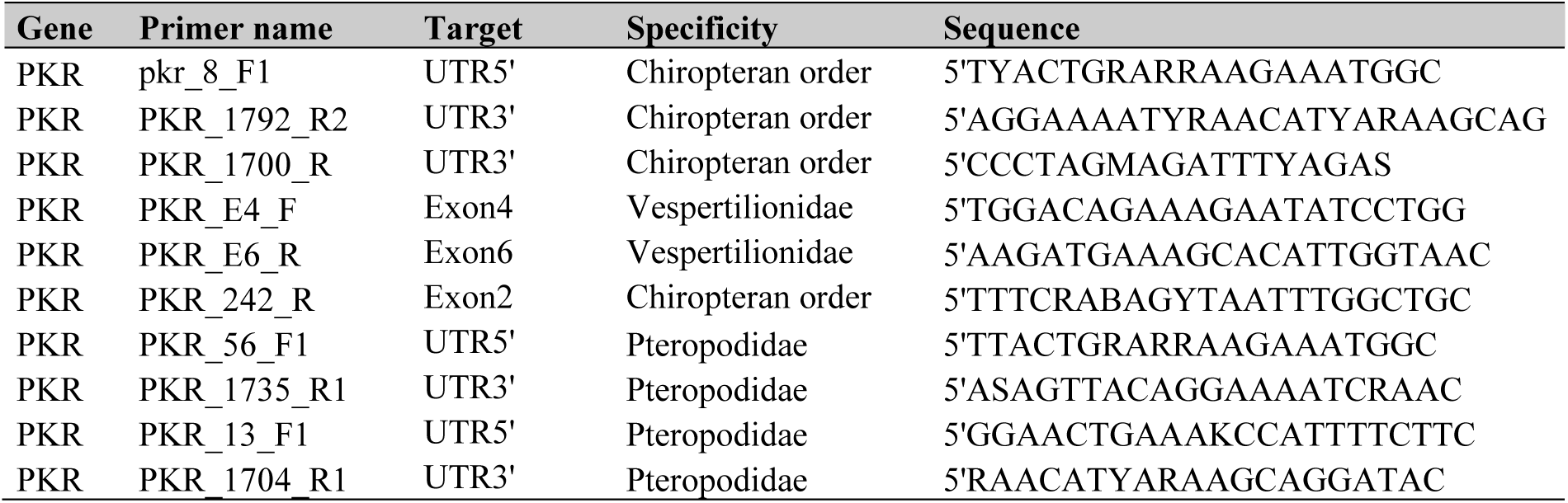
Primer sets used in the study. The primers listed here are universal to the chiropteran order or to specific families.

### Supplementary Figures

**Figure S1.**
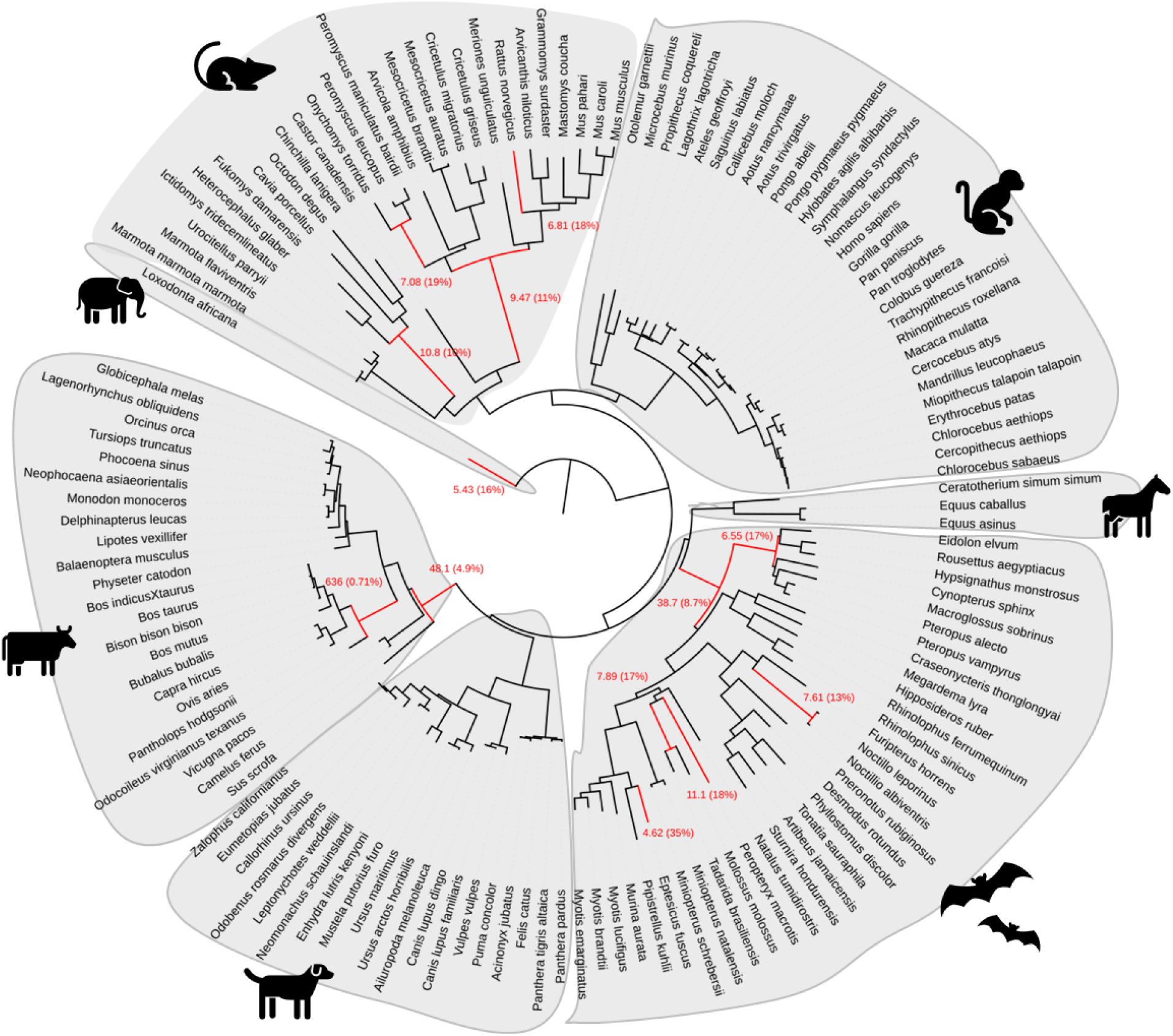
PKR has evolved under episodic positive selection across mammals. Maximum likelihood phylogenetic tree of mammalian PKR showing the branches under significant positive selection (p-value <0.05, in red) in artiodactyls, carnivores, bats, perissodactyls, primates, rodents and proboscideans. Analysis was performed using aBSREL from the HYPHY package. The numbers in brackets indicate the estimated values of the ω at the branch. The scale bar indicates The scale bar indicates the number of substitutions per site.

**Figure S2.**
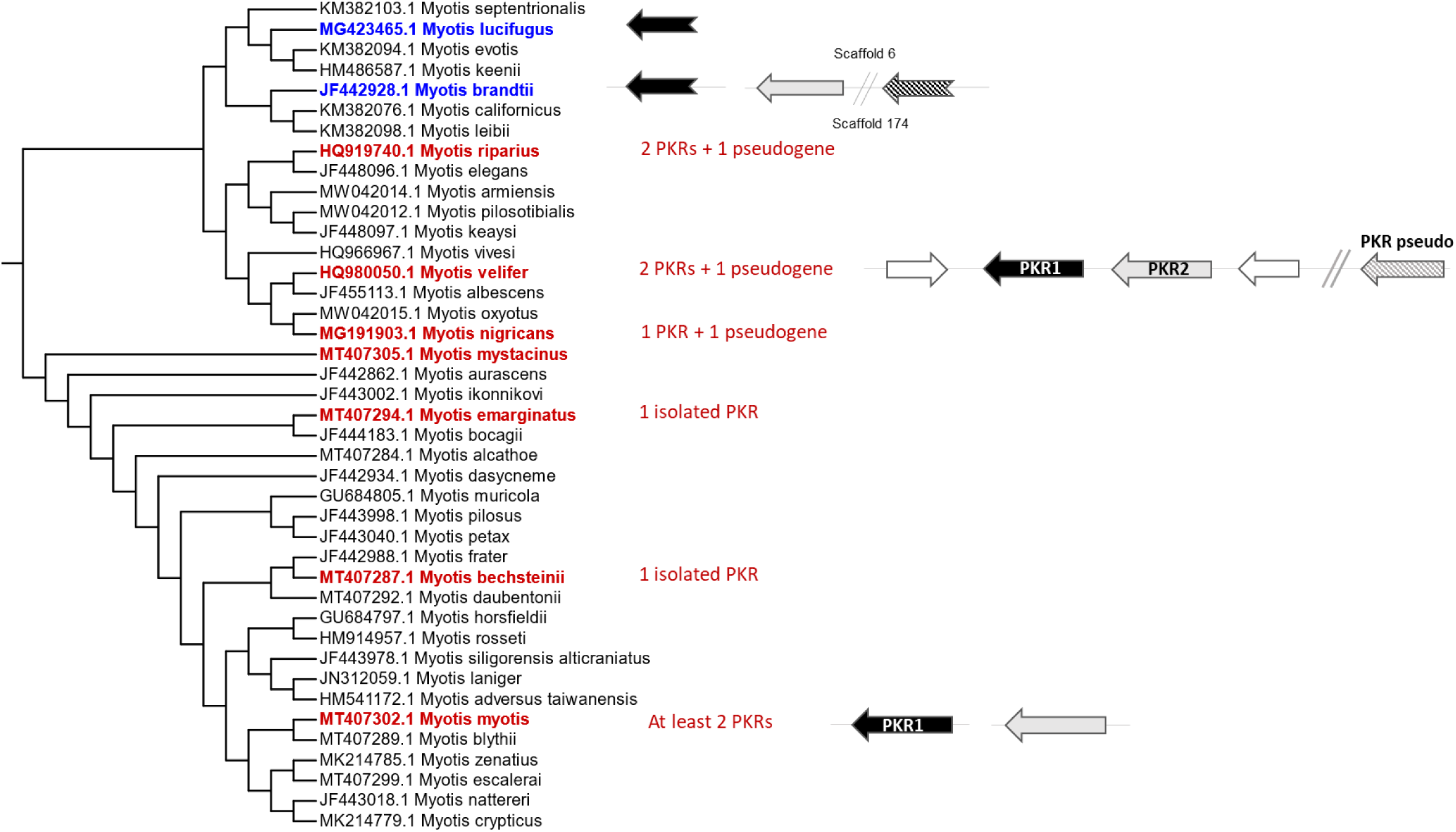
Intra-species characterization of PKR duplication in *Myotis* species. In the left, a cladogram representing the Myotis phylogenetic tree, based on the Cytochrome oxidase I gene. Sequences were retrieved from GenBank and aligned with Prank software. The PhyML program was used to build the phylogenetic tree, using the best fitting substitution model (TN93+R) – inferred by the SMS program. The Myotis species that were analyzed in this study are highlighted. In blue are the publicly available genomes, and in red, are the newly sampled species in this study. For each analyzed species, the number of PKR copies fully or partially isolated by PCR – cloning method are indicated.

**Figure S3.**
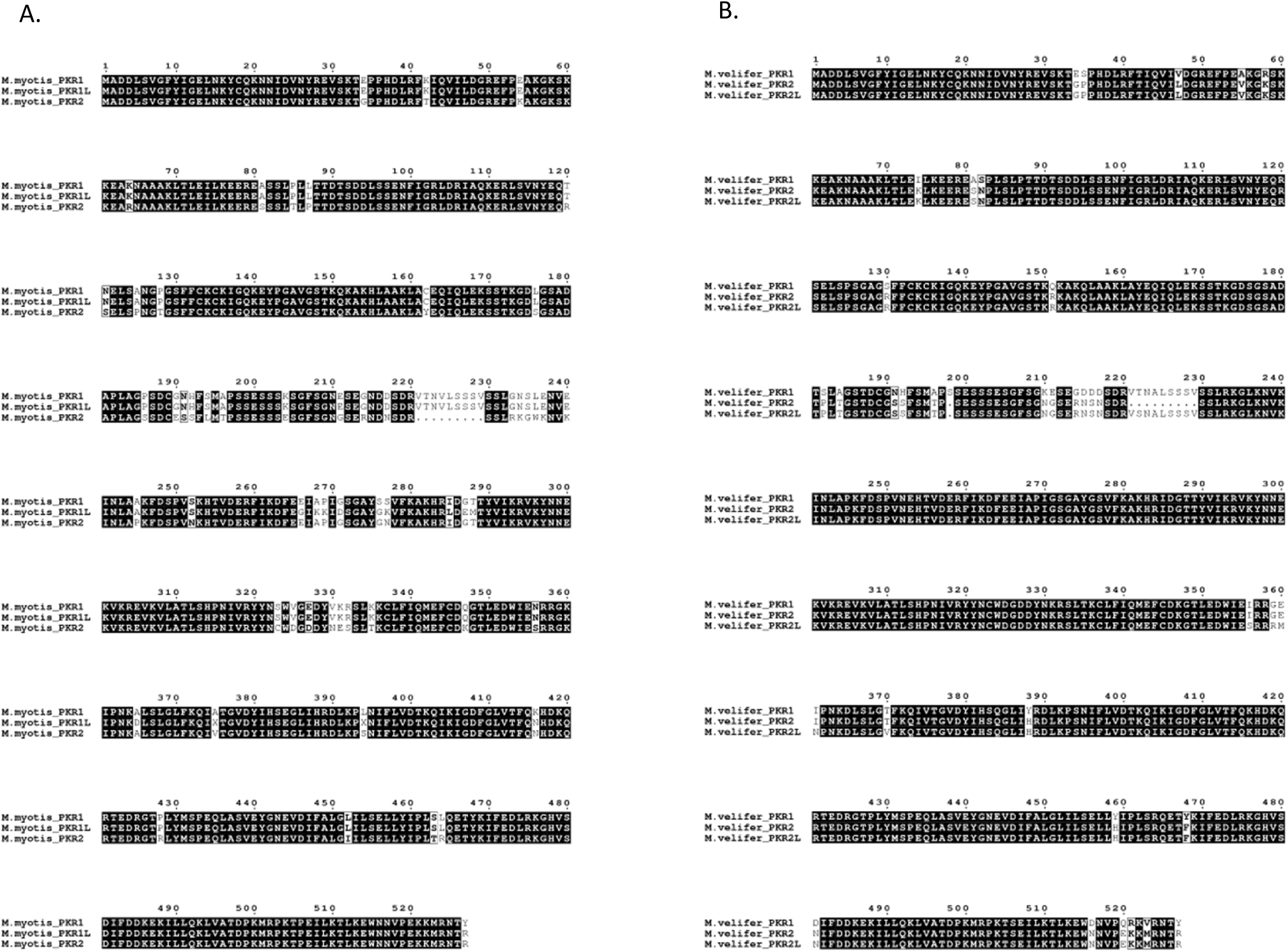
Protein alignment of *M. myotis* and *M. velifer* PKR paralogs. Visualization was obtained with ESPript^94^

**Figure S4.**
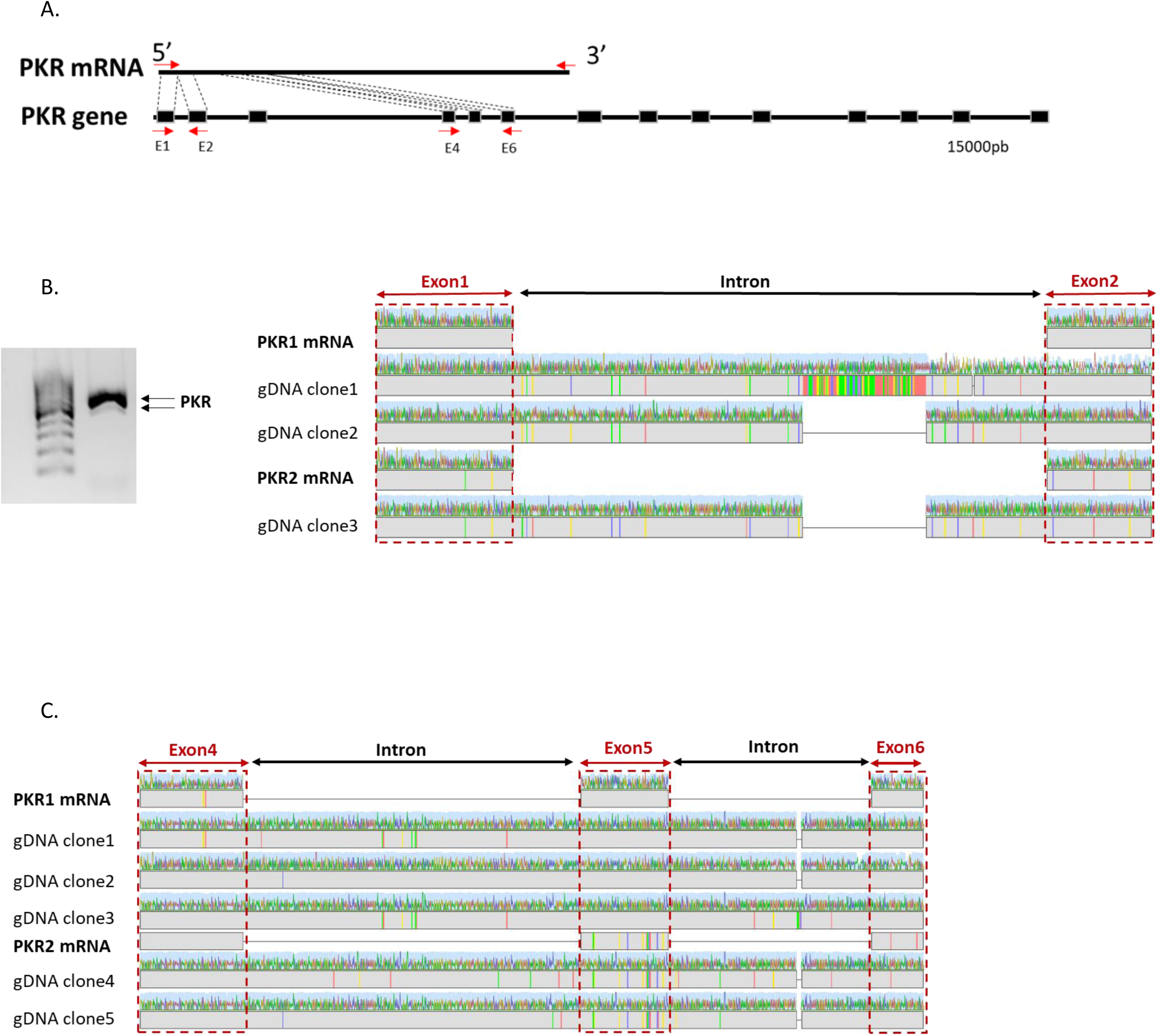
Characterization of PKR duplication in *Myotis myotis.* PCR amplification of different regions (E1-E2, and E4-E6) of *EIF2AK2* gene from *M. myotis*. **A**. Schematic representation of the PCR strategy. Red arrows represent the primers, black boxes are exons and internal bars are introns. **B-C**. electrophoresis migration of PCR products (left) and electropherogram alignment of isolated variants (right) obtained with specific primers of exons 1 and 2 (B) and exons 4 and 6 (C).

**Figure S5.**
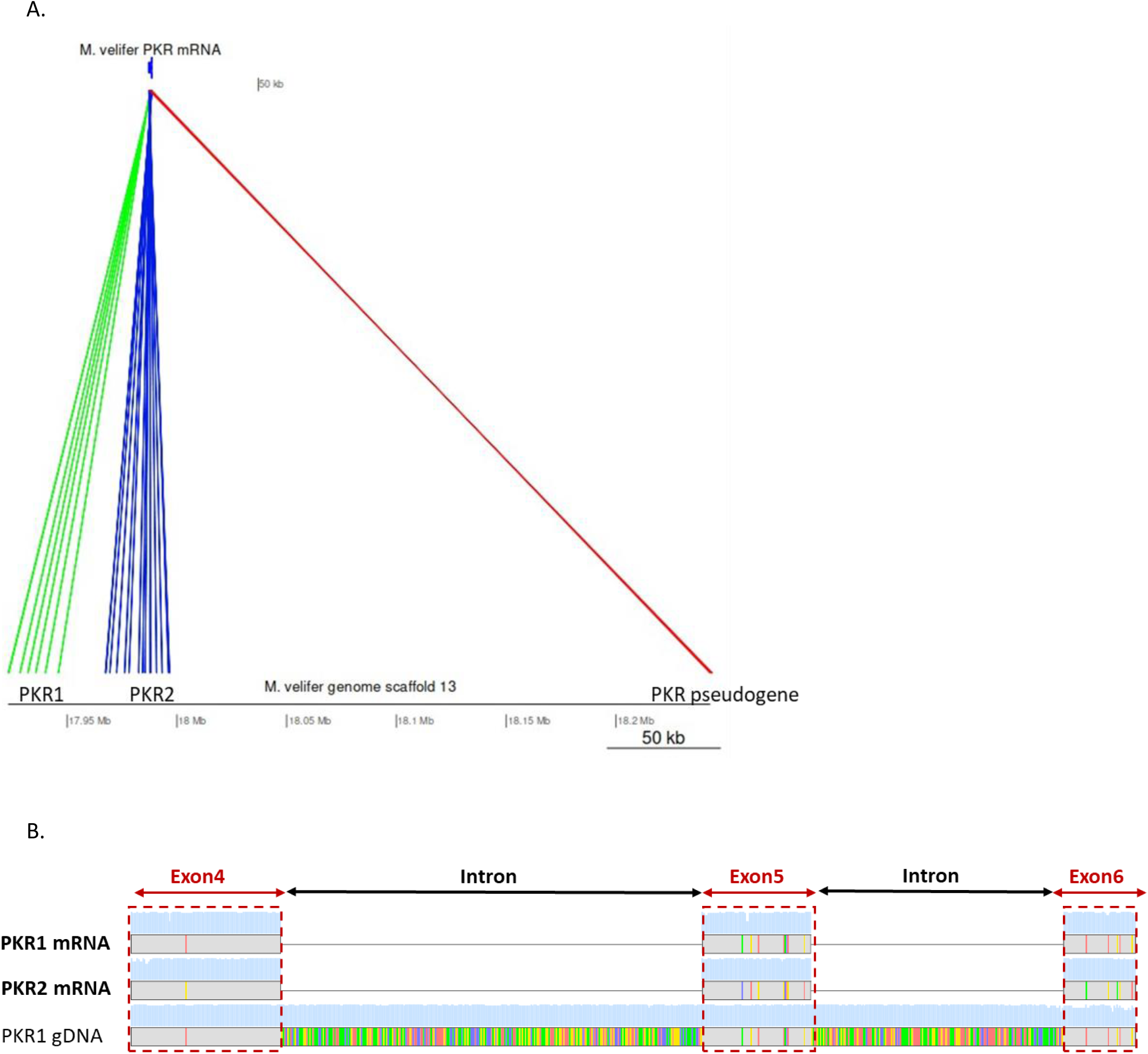
Additional characterization of PKR duplicates in *Myotis velifer*. **A.** Tblastn search of PKR in *M. velifer* genome indicating the presence of three copies in the same scaffold (black bar on the bottom). PKR mRNA sequence from M. velifer was used as a query (blue arrow on top), with a cut-off of 10e^-05^. **B-C**. Electropherogram alignment of *EIF2AK2* sequence exons 1 and 2 (B) and exons 4 and 6 (C) isolated from *M. velifer* genomic DNA using specific primers.

**Figure S6.**
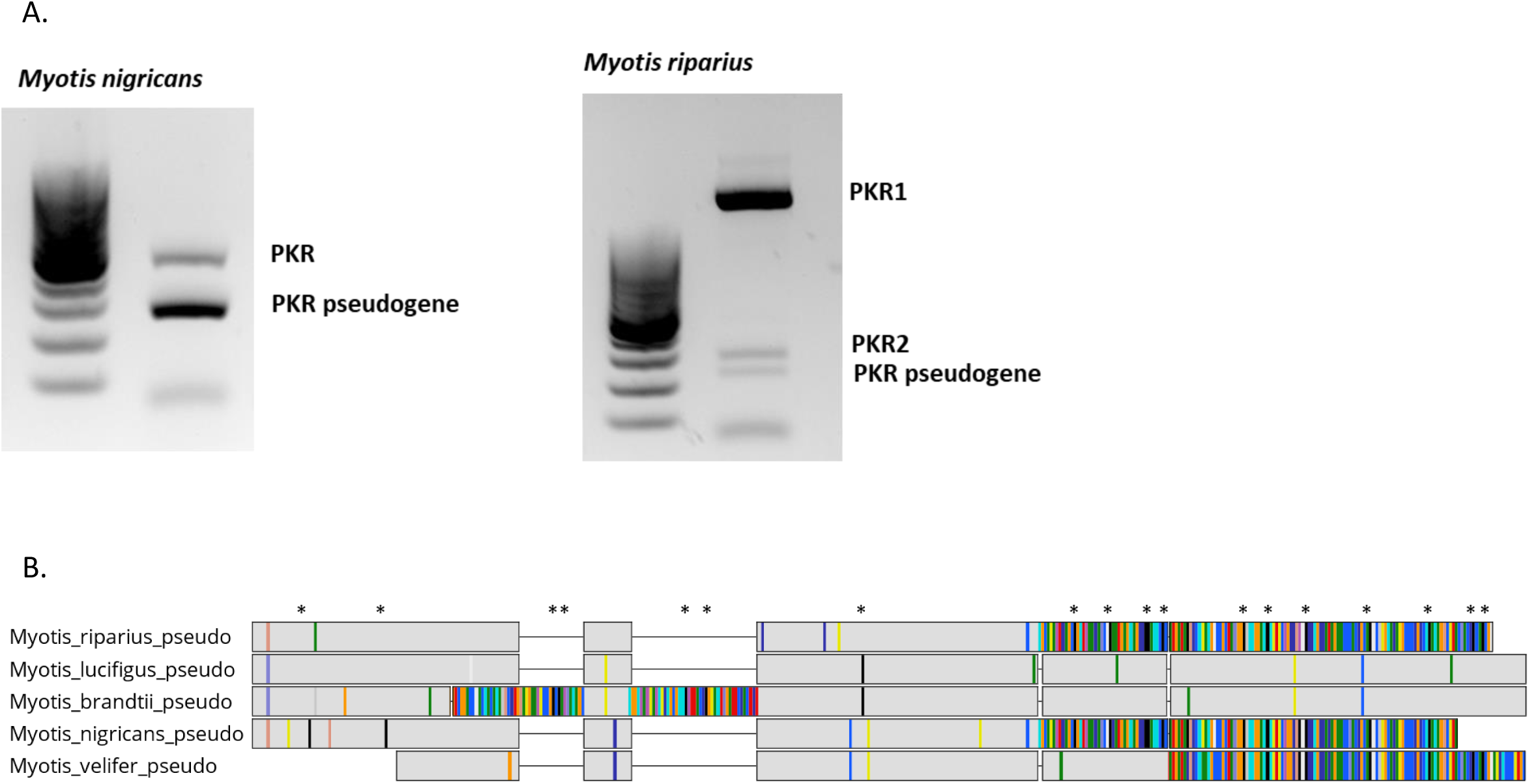
Genomic evidences of PKR duplication and pseudogeneization in New World *Myotis* species. **A**. PCR amplification of exons 4 to 6, revealing the existence of *EIF2AK2* pseudogene in *M. velifer* and *M. riparius*. The electrophoresis migrations of PCR products obtained with specific primers of exons 4 and 6 are shown. **B**. Sequence alignment of isolated PKR pseudogenes, with the presence of multiple stop codons (*).

**Figure S7.**
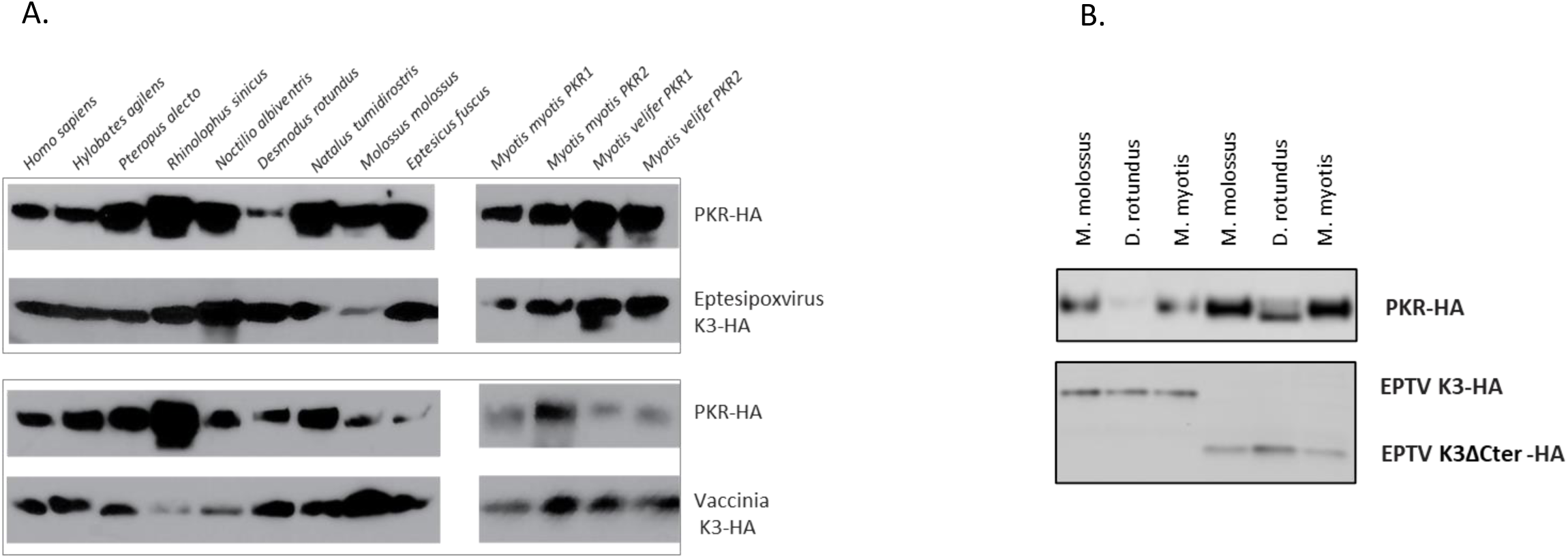
Expression of bat PKRs, as well as EPTV and VACV K3s in yeast spot assays. **A.** PKR ortholog co-expression with EPTV and VACV K3. **B.** PKR ortholog co-expression with EPTV K3 or EPTV K3 mutant.

**Figure S8.**
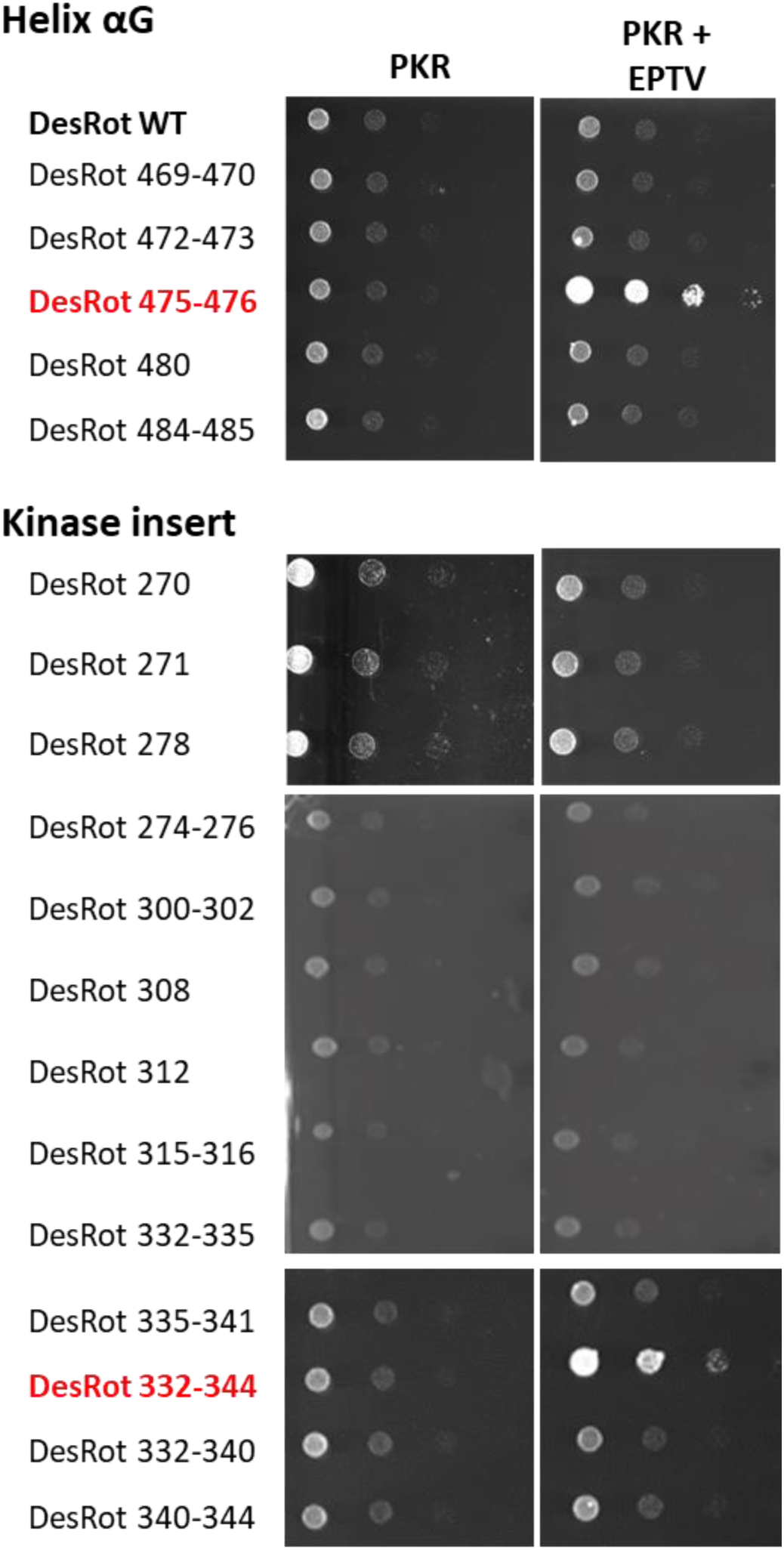
PKR mutants identifying the genetic determinants of PKR susceptibility to eptesipox virus K3. Spot assays of *D. rotundus* PKR mutants identifying the genetic determinants of PKR susceptibility to eptesipox virus and variola K3 antagonism. The Helix αG and kinase insert mutants were generated by site-directed mutagenesis of *D. rotundus* PKR, by swapping the corresponding residues from *M. myotis* PKR2.

**Figure S9.**
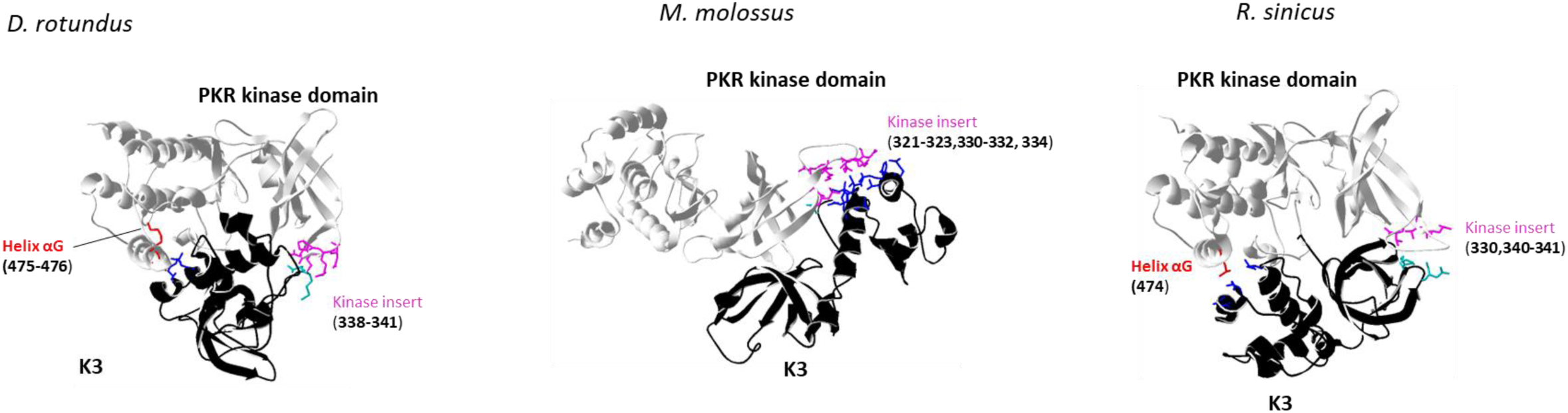
Protein-protein complex structure between PKR and EPTV K3. Protein-protein complex structure between epetsipoxvirus K3 and *D. rotundus*, or *M. molossus* PKRs inferred by Hdock software. The residues involved in the K3-PKR interface are colored in red for PKR Helix αG and magenta for kinase insert . In eptesipox virus K3, C-terminal insertion is colored in dark bue, and other contact residues are in light blue. The docking model shows, different contact affinity depending on the PKR structure (related to yeast functional assay, Figure 2).

**Figure S10.**
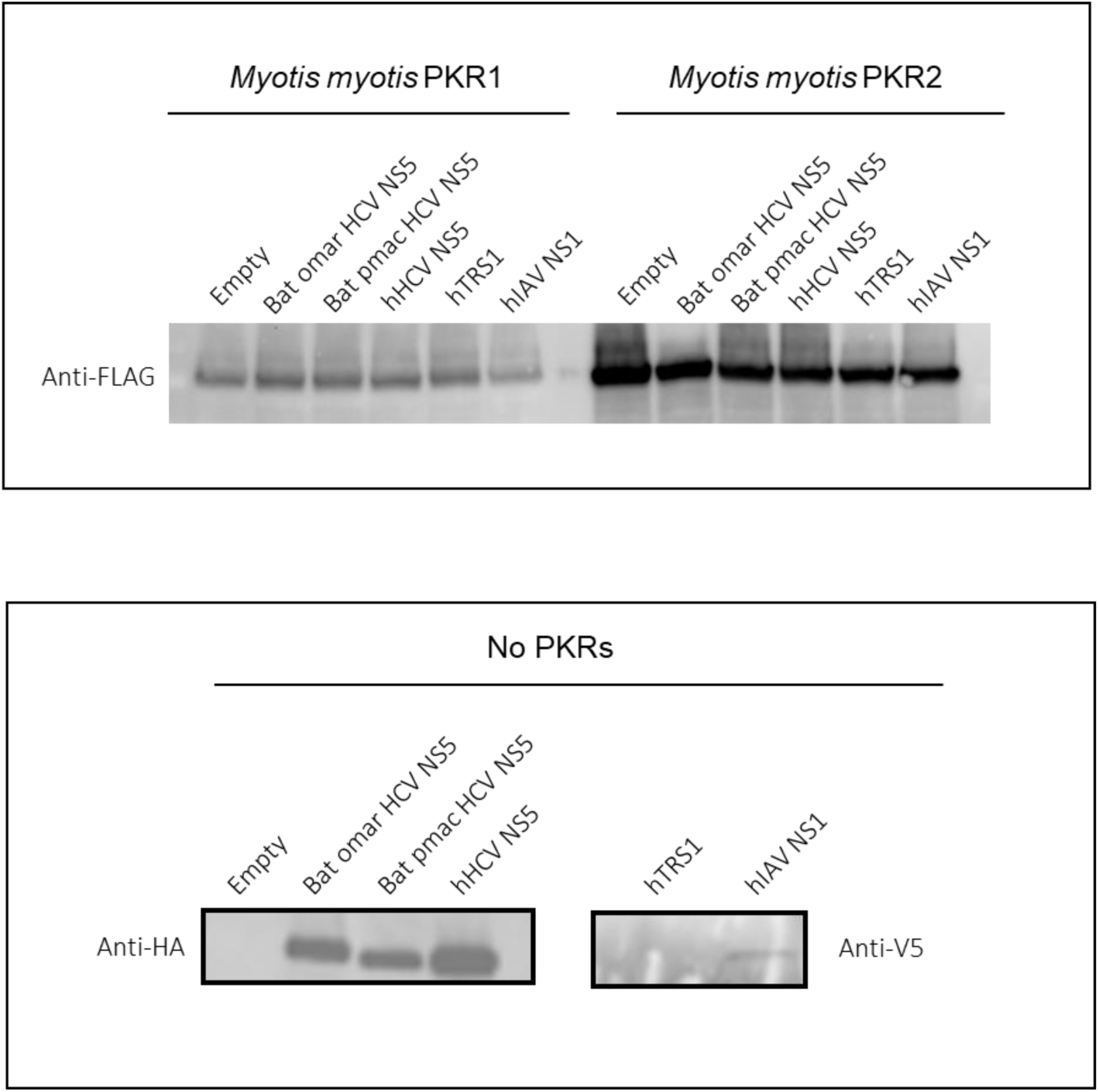
Immunoblot assaying the expression of bat PKRs and the viral antagonists, NS5A, NS1 and TRS1, in luciferase assays. Despite a two-fold difference between PKR1 and PKR2 (estimated with ImageLab software; BioRad), PKR1 showed similar levels of translation inhibition (Figure 2).

**Figure S11.**
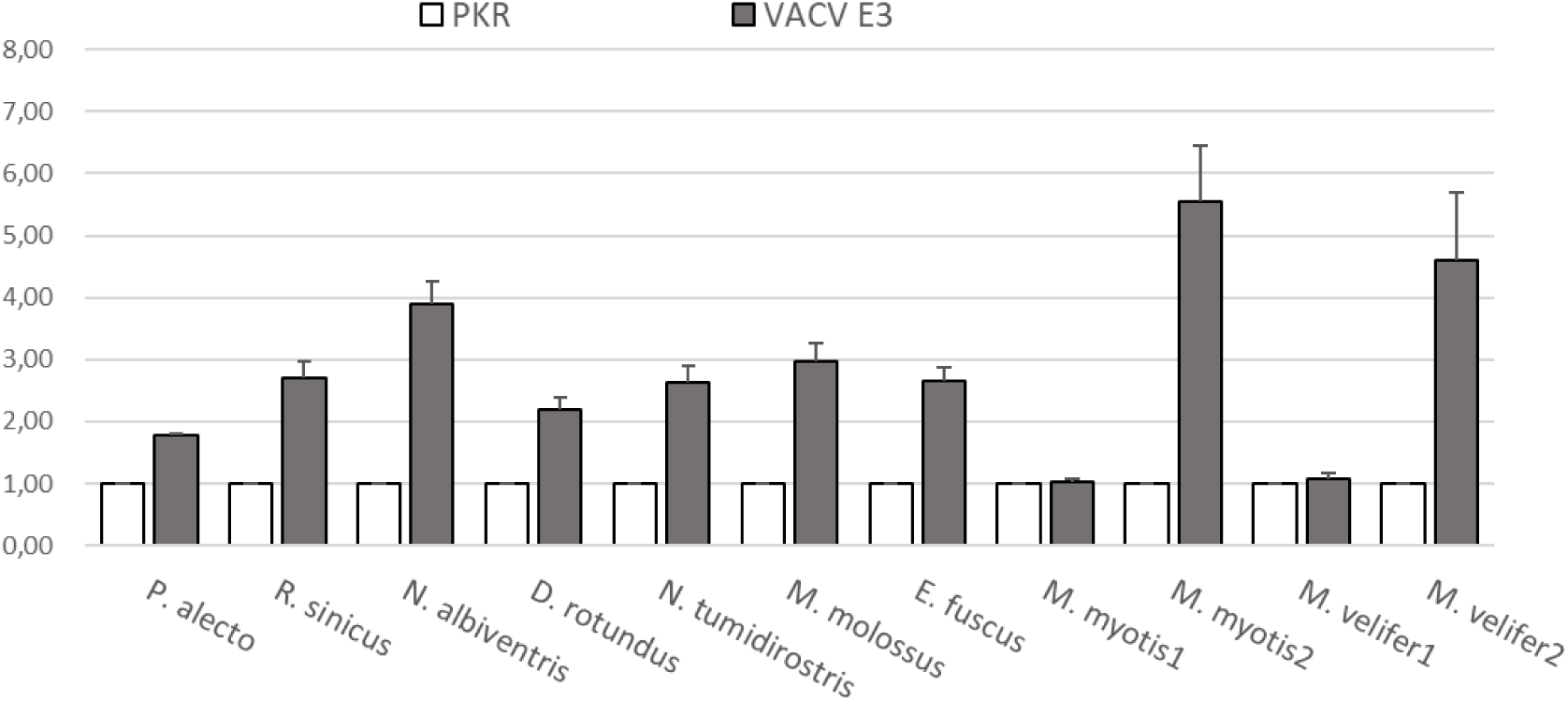
Luciferase assay showing species-specific sensitivity of PKRs to VACV E3 antagonism. All bat PKRs variants showed significant level of susceptibility to VACV E3. Three independent experiments were conducted; values represent the mean of the triplicates. Luciferase activity was normalized to the control condition in which cells were transfected with PKR and empty vector (x axis). Error bars indicate SEM

